# IDH-mutant inhibitors enhance the sensitivity of IDH1-mutant gliomas to cysteine-methionine deprivation and ferroptosis

**DOI:** 10.64898/2026.01.30.702627

**Authors:** Angeliki Mela, Abby Brand, Aayushi Mahajan, Athanassios Dovas, Nelson Humala, Smriti Kanangat, Allison C Kleinstein, Sandra Leskinen, Trang TT Nguyen, Qiuqiang Gao, Pavan S Upadhyayula, Jia Guo, Brian Gill, Markus D Siegelin, Peter A Sims, Brent R Stockwell, Jeffrey N Bruce, Peter Canoll

## Abstract

Cysteine is essential for synthesizing glutathione, the brain’s main antioxidant, and cysteine deprivation can trigger ferroptosis. Here, using a new mouse model of IDH1-mutant glioma that recapitulates the characteristics of human IDH1-mutant low-grade gliomas, we demonstrate that IDH1-mutant glioma cells are significantly more vulnerable to cysteine deprivation alone or in combination with the ferroptosis inducer RSL3, compared to IDH1-wildtype glioma cells. In addition, treatments with the IDH-mutant inhibitors vorasidenib and ivosidenib further sensitize the cells to ferroptosis. Metabolomics analysis reveals that IDH1-mutant cells have altered cysteine and methionine metabolism with deficiency in transsulfuration, which is further exacerbated by cysteine-methionine deprivation and IDH-mutant inhibitors. Furthermore, dietary cysteine-methionine deprivation alone or in combination with convection-enhanced delivery of RSL3 or ivosidenib *in vivo* significantly prolongs survival of IDH1-mutant tumor-bearing mice. Our findings suggest that targeting cysteine and methionine metabolism in combination with IDH-mutant inhibition provides promising therapeutic strategies for IDH1-mutant gliomas.

## Introduction

Mutations in isocitrate dehydrogenase 1 (IDH1) define a distinct subset of diffuse gliomas characterized by slower progression, diffuse infiltration, and profound metabolic reprogramming^1,2,3^. The neomorphic activity of mutant-IDH1 converts α-ketoglutarate (α-KG) to the oncometabolite D-2-hydroxyglutarate (2-HG), which inhibits α-KG–dependent dioxygenases and perturbs redox homeostasis and intermediary metabolism^4,5,6,7,8,9^. Although the epigenetic and signaling consequences of 2-HG accumulation are well characterized, the broader metabolic consequences of mutant-IDH (IDHmut) and their functional implications for tumor growth and therapeutic response remain incompletely defined. Furthermore, the metabolic effects of IDHmut-inhibitors currently used in the treatment of low-grade IDH-mutant gliomas have not yet been fully elucidated^10,11,12^.

Emerging studies indicate that IDH1-mutant gliomas exhibit a chronic state of oxidative and metabolic stress^13,14^. Among these changes, alterations in cysteine and methionine metabolism are important, yet understudied features of the IDH1-mutant metabolic landscape^13^. These pathways play essential roles in maintaining glutathione synthesis, methylation potential, and redox buffering^15,16^. Dysregulation of transsulfuration and methionine cycle may render IDH1-mutant cells dependent on exogenous cysteine and vulnerable to fluctuations in extracellular nutrient availability^17,18,19^.

Recent work suggests that alterations in redox state may sensitize cells to ferroptosis, a form of regulated cell death driven by lipid peroxidation and glutathione depletion^20,21,22,23^. By constraining cysteine and methionine metabolism, mutant-IDH1 may create a permissive context for ferroptotic vulnerability—linking amino acid metabolism, oxidative stress, and cell death sensitivity. Here, we demonstrate that the IDH1 mutation alters cysteine and methionine metabolism to impose a redox constraint that sensitizes glioma cells to ferroptosis, revealing a previously unrecognized metabolic liability with therapeutic potential, that can be further exploited by combining cysteine-methionine deprivation with IDHmut-inhibitors.

## Results

### IDH1(R132H) significantly delays glioma progression

To generate an IDH1-mutant glioma model, we performed stereotactic intracranial injections of PDGFA-IRES-CRE expressing retrovirus into the subventricular zone of neonatal (p4) Idh1(R132H)^KI^/+; p53^fl/fl^; RiboTag (IDH1-mutant) and +/+; p53^fl/fl^; RiboTag (IDH1-wildtype) mice (Figure 1A). We monitored tumor progression using MRI (Figure 1B). Our results show that mice of both genotypes formed tumors, however the IDH1 mutation significantly delays glioma progression (Figure 1B, C). Specifically, the IDH1-mutant, PDGFA-overexpressing, p53-deleted mice start developing lesions visible by MRI at 70-190 days post injection (dpi), while IDH1-wildtype mice reach end stage at 50-70 dpi (Figure 1C, D). Histopathological analysis of end-stage primary IDH1-mutant tumors revealed diffuse brain infiltration and hypercellularity, but in contrast to IDH1-wildtype tumors, showed few mitotic figures and an absence of vascular proliferation or necrosis (Figure 1C, Supplementary Figure 1). Immunohistochemistry for HA, the RiboTag reporter, labeled retrovirally infected tumor cells in both the IDH1-mutant and IDH1-wildtype models. In both models, tumor cells robustly expressed Olig2 and displayed a diffusely infiltrative growth pattern, intermingling with GFAP⁺ astrocytes, NeuN⁺ neurons, and Iba1⁺ microglia. Notably, these non-neoplastic cell populations were more prominent in the IDH1-mutant model (Figure 1C). A major distinction between the two models is their Ki67 proliferation index. The IDH1-wildtype mouse model exhibited a Ki67 index of 55.35% at 58 dpi (end stage), whereas the IDH1-mutant model showed markedly lower indices of 5.9% at 60 dpi and 15.45% at 250 dpi (Figure 1E), in line with the diffusely infiltrative, low-grade characteristics of human IDH1-mutant gliomas.

**Figure 1.**
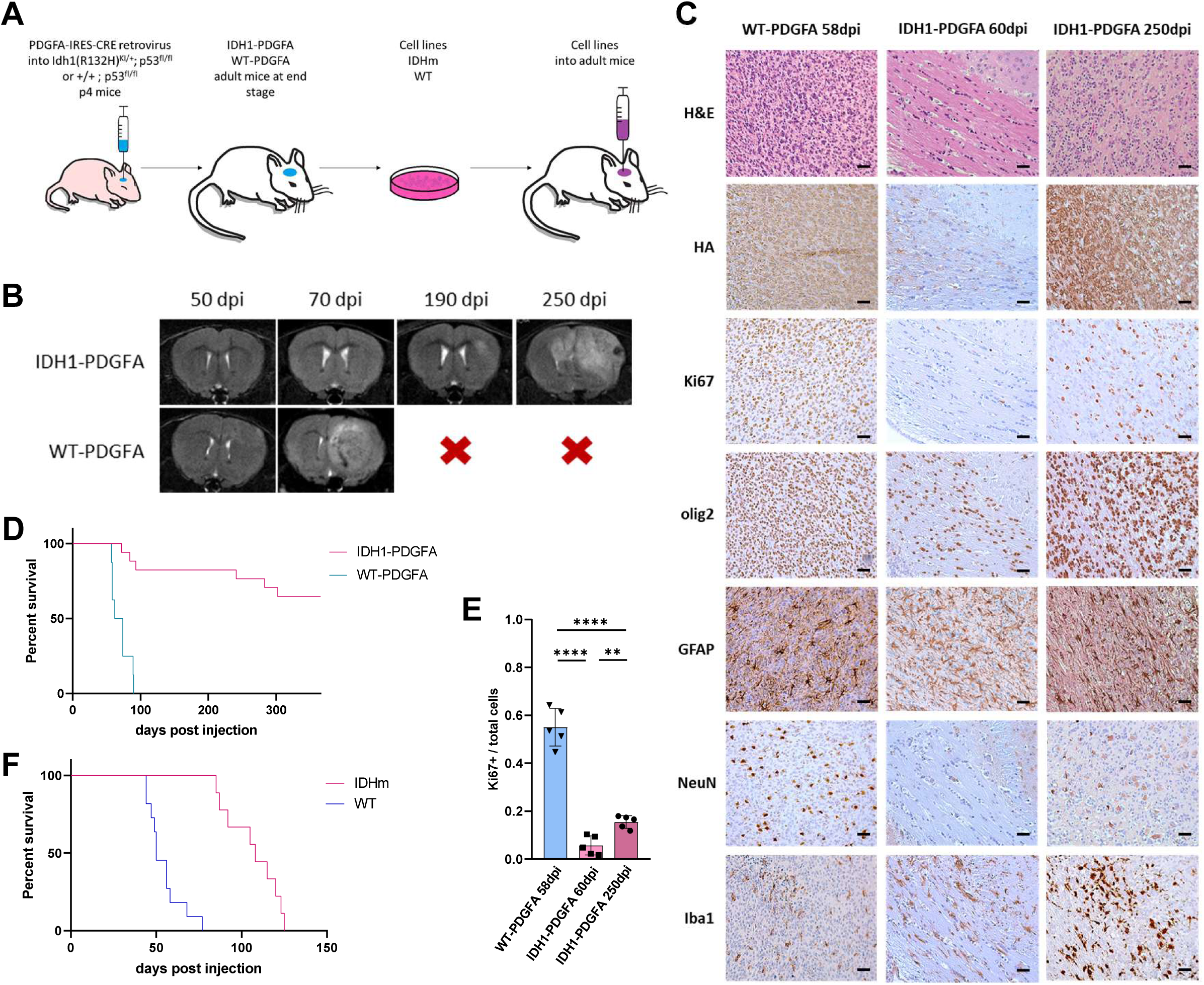
IDH1(R132H) mutation significantly delays glioma progression. **A.** Diagram of experimental design of stereotactic intracranial injections of PDGFA-IRES-CRE retrovirus in Idh1(R132H)^KI/+^; p53^fl/fl^; RiboTag and +/+; p53^fl/fl^; RiboTag neonatal (p4) mice, cell line generation and orthotopic injections. **B.** T2 MRI images showing glioma progression at 50, 70, 190 and 250 dpi in IDH1-mutant (IDH1-PDGFA) and IDH1-wildtype (WT-PDGFA) mice. **C.** Immunohistochemistry of a WT-PDGFA tumor at 58dpi (end stage), an IDH1-PDGFA tumor at 60dpi (early stage) and an IDH1-PDGFA tumor at 250dpi (end stage) for HA, Ki67, olig2, GFAP, NeuN and Iba1. Anti-HA labels retrovirus-infected cells, hematoxylin was used as nuclear stain. Bar=100mm. **D.** Survival curves after retrovirus injections in neonatal (p4) mice. Median survival: IDH1-PDGFA undefined, WT-PDGFA 68dpi. Log-Rank (Mantel-Cox) test: p<0.0001****. **E.** Ki67 labeling index in WT-PDGFA 58dpi, IDH1-PDGFA 60dpi and IDH1-PDGFA 250dpi tumors. Ki67+ cells divided by total (hematoxylin positive) cells counted in 3 mice per group, two 20x fields per mouse (WT-PDGFA 58dpi: 3820 Ki67+ / 6901 total, IDH1-PDGFA 60dpi: 172 Ki67+ / 2914 total, IDH1-PDGFA 250dpi: 768 Ki67+ / 4969 total). t-test WT-PDGFA 58dpi vs IDH1-PDGFA 250dpi p=0.000005****, WT-PDGFA 58dpi vs IDH1-PDGFA 60dpi p=0.000002****, IDH1-PDGFA 60dpi vs IDH1-PDGFA 250dpi p=0.001772**. **F.** Survival curves after orthotopic injections of IDH1-mutant (IDHm) and IDH1-wildtype (WT) cells into adult B6 mice. Median survival: IDHm 108dpi, WT 50dpi. Log-Rank (Mantel-Cox) test: p<0.0001****.

At end stage, we harvested tumor cells and generated primary cell cultures from IDH1-mutant and IDH1-wildtype tumors. To confirm tumorigenic potential of these cell lines, we performed stereotactic intracranial injections into adult mice (Figure 1A, F). All mice injected with either IDH1-mutant or IDH1-wildtype cell lines form tumors, significantly faster than the primary tumor model that each cell line was generated from. Notably, mice injected with IDH1-mutant cells survive significantly longer than mice injected with IDH1-wildtype cells (Figure 1F). IDH1-mutant cells produce high levels of 2-HG both *in vitro* and *in vivo*, as detected by MEGA-PRESS MRS (Supplementary Figure 2A). MEGA-PRESS MRS also detects lower glutathione levels and higher glutamate to GABA ratio in IDH1-mutant mouse tumors *in vivo* (Supplementary Figure 2B, C), consistent with human IDH1-mutant gliomas^24,25^. These results suggest that our mouse model recapitulates key characteristics of human IDH1-mutant gliomas and can be used for further *in vitro* and *in vivo* studies.

### IDH1-mutant glioma cells are sensitive to cysteine-methionine deprivation and ferroptosis in vitro

Previously we have shown that cysteine-methionine deprivation (CMD) sensitizes IDH-wildtype glioma cells to ferroptosis^23^. To test this in our IDH1-mutant glioma cells we used RSL3, a GPX-4 inhibitor. We treated both IDH1-mutant and IDH1-wildtype cell lines with different concentrations of RSL3 for 17 hours with or without cysteine and methionine. Our results reveal that IDH1-mutant cells are significantly more sensitive to RSL3-induced ferroptosis in both control and CMD conditions (Figure 2A, B). CMD enhances sensitivity to RSL3, while treatment with the ferroptosis inhibitor Ferrostatin-1 prevents cell death in all conditions. These results were also confirmed by showing RSL3-mediated induction of lipid peroxidation as evidenced by green fluorescence shift in Bodipy-C11, following the addition of RSL3 alone or in combination with CMD (Supplementary Figure 3A). Our results show that IDH1-mutant cells are significantly more sensitive to RSL3 treatment compared to IDH1-wildtype cells and CMD significantly enhances their sensitivity. We also investigated the expression of Slc7a11 and Atf4, which are both transcriptional hallmarks of cellular response to ferroptosis^26,27,28^. RT-qPCR showed that there are significant increases of Slc7a11 and Atf4 transcripts in both IDH1-mutant and IDH1-wildtype cells after 17-hour treatment with CMD alone or with RSL3 (Supplementary Figure 3B).

**Figure 2.**
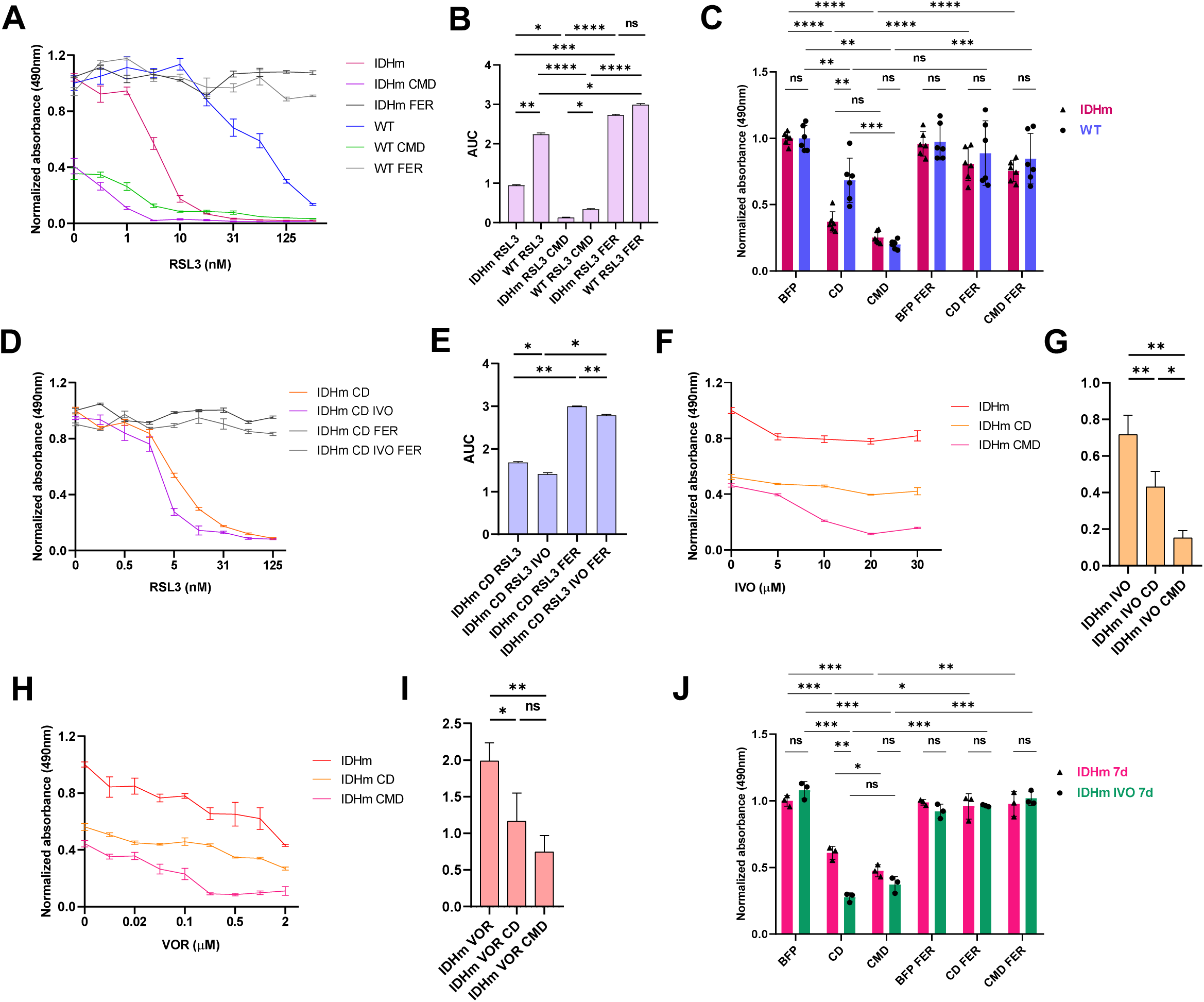
Cysteine-methionine deprivation and IDHmut-inhibitors enhance the sensitivity of IDH1-mutant cells to ferroptosis in vitro. **A.** Dose response curves after 17-hour treatment of IDH1-mutant (IDHm) and IDH1-wildtype (WT) cells with RSL3 (0, 0.5, 1, 5, 10, 15, 31, 62, 125, 250 nM) alone, with CMD or 2mM Ferrostatin-1 (FER) (IC50 WT 52.6, IDHm 5.1, WT-CMD 4.513, IDHm-CMD 0.3573). **B.** Area under the curve (AUC) plotted from three independent experiments as in A. **C.** MTS viability assay after 24 hours in control (BFP), CD or CMD conditions in the presence or absence of 2mM Ferrostatin-1. **D.** Dose response curves of IDH1-mutant cells after 17-hour treatments with RSL3 (0, 0.1, 0.5, 1, 5, 10, 31, 62, 125 nM) in CD conditions with or without 20mM IVO or 2mM Ferrostatin-1 (IC50 IDHm-CD 5.832, IDHm-CD-IVO 2.106). **E.** Area under the curve (AUC) plotted from three independent experiments as in D. **F.** Dose response curves after 17-hour treatment of IDH1-mutant cells with IVO (0, 5, 10, 20, 30 mM) in control, CD or CMD conditions (IC50 IDHm-CMD 9.526). **G.** Area under the curve (AUC) plotted from three independent experiments as in F. **H.** Dose response curves after 17-hour treatment of IDH1-mutant cells with VOR (0, 0.005, 0.02, 0.05, 0.1, 0.2, 0.5, 1, 2 mM) in control, CD or CMD conditions (IC50 IDHm 0.9281, IDHm-CD 0.7768, IDHm-CMD 0.0677). **I.** Area under the curve (AUC) from three independent experiments as in H. **J.** MTS viability assay of IDH1-mutant cells grown for 7 days in the absence or presence of 20mM IVO and subsequently treated in control (BFP), CD or CMD conditions for 17 hours with or without 2mM Ferrostatin-1 (FER). Data plotted as mean ± SD. Statistics assessed using t-test. Significance denoted by: *p < 0.05, **p < 0.01, ***p < 0.001, ****p<0.0001, ns: not significant.

To test whether IDH1-mutant cells are more sensitive to cysteine deprivation alone, we treated both IDH1-mutant and IDH1-wildtype cells with media made without cysteine (CD), or without cysteine and methionine (CMD) for 24 hours (Figure 2C). While both cell lines are sensitive to CMD, IDH1-mutant cells are significantly more sensitive than IDH1-wildtype cells to cysteine depletion alone. Both effects were rescued by Ferrostatin-1. To further investigate this sensitivity to cysteine deprivation, we used another ferroptosis inducer, IKE, that blocks cysteine uptake via the Slc7a11/Slc3A2 System Xc^-^ cysteine-glutamate antiporter. We treated IDH1-mutant and IDH1-wildtype cells with different concentrations of IKE in control media (containing normal levels of cysteine and methionine) for 17 hours. Our results show that IDH1-mutant cells are significantly more sensitive to IKE-induced ferroptosis, which is rescued by Ferrostatin-1 (Supplementary Figure 3C, D). Furthermore, we treated IDH1-mutant and IDH1-wildtype cells with RSL3 in cysteine-deprived (CD) media. Our results show that IDH1-mutant cells are significantly more sensitive to CD and ferroptosis compared to IDH1-wildtype cells and these effects are rescued by Ferrostatin-1 (Figure 2D, E; Supplementary Figure 4A, B).

### IDHmut-inhibitors enhance the sensitivity to cysteine-methionine deprivation in vitro

The development of IDHmut-inhibitors and their use as promising therapeutic glioma treatments prompted us to test potential treatment effects in our IDH1-mutant mouse model as well as possible synergistic or antagonistic effects with CMD and ferroptosis inducers. We tested two FDA-approved IDHmut-inhibitors, ivosidenib (IVO) which selectively inhibits mutant IDH1, and vorasidenib (VOR), a dual mutant IDH1/2 inhibitor^10,11,12,29^.

To assess the effects of IDHmut-inhibitors on cell viability, we treated IDH1-mutant and IDH1-wildtype cells *in vitro* with ivosidenib (IVO) or vorasidenib (VOR) under control, CD, or CMD conditions for 17 hours (Figure 2F–I). IVO has minimal effect on cell viability of IDH1-wildtype cells across growth conditions (Supplementary Figure 4C, D). In contrast, IVO induced dose-dependent reductions in viability of IDH1-mutant cells specifically when grown in CMD and to a lesser extent in CD conditions, but not in control media (Figure 2F, G). VOR similarly showed minimal effects on IDH1-wildtype cell viability at concentrations below 1 µM (Supplementary Figure 4E, F). However, VOR caused dose-dependent decreases in viability in IDH1-mutant cells at concentrations below 1 µM even in control media, and these effects were significantly enhanced under CD and CMD conditions (Figure 2H, I). At concentrations above 0.05 µM, VOR reduced IDH1-mutant cell viability by approximately 15% in control conditions, ∼50% under CD conditions, and ∼70% under CMD conditions (Figure 2H). Although these differences between IVO and VOR may reflect differences in potency or the dual targeting of IDH1 and IDH2 by VOR^29^, both inhibitors markedly reduced IDH1-mutant cell viability when combined with CMD. Together, these findings indicate that cysteine-methionine deprivation sensitizes IDH1-mutant glioma cells to IDH-mutant inhibition, irrespective of the specific inhibitor used.

To investigate whether IDHmut-inhibitors affect the expression of Slc7a11 in our IDH1-mutant glioma cells, we treated IDH1-mutant cells with 20mM IVO or 100nM VOR for 24 and 48 hours and subsequently performed qRT-PCR for Slc7a11 (Supplementary Figure 4G). Both inhibitors significantly increase Slc7a11 transcripts after 24 hours, and IVO continues to do so at 48 hours. These results suggest that combining IDHmut-inhibitors with CMD and ferroptosis inducers may cause an increased dependence on extracellular cysteine and induce ferroptotic cell death.

To further investigate this, we treated IDH1-mutant and IDH1-wildtype cells in CD (normal methionine) conditions with 20mM IVO and various doses of RSL3 (Figure 2D, E; Supplementary Figure 4A, B). Our results show that IDH1-mutant cells are significantly more responsive to triple combination treatment of RSL3 with IVO and CD compared to just RSL3 and CD. In contrast, IDH1-wildtype cells are not significantly affected by the addition of IVO or cysteine deprivation. To assess the effects of more prolonged treatment with IDHmut-inhibitors *in vitro*, IDH1-mutant cells were grown for 7 days with or without 20mM IVO and subsequently treated with cysteine-deprived or cysteine-methionine deprived media (CD or CMD) (Figure 2J). Our results show that IDH1-mutant cells treated for 7 days with IVO are more sensitive to CD than non-treated cells. CMD *in vitro* has the same robust effect in both IVO-treated and non-treated cells.

### Metabolomic analyses reveal that IDH1-mutant glioma cells have transsulfuration deficiency that is exacerbated by cysteine-methionine deprivation and IDHmut-inhibitors

To further explore these metabolic vulnerabilities, we performed targeted metabolite profiling for more than 200 metabolites on our IDH1-mutant and IDH1-wildtype mouse cell lines grown under different conditions. Under control conditions, IDH-mutant cells showed significantly higher levels of 2-HG along with significant alterations in multiple additional metabolites compared to IDH-wildtype cells (Figure 3A, B). Pathway analysis showed that cysteine and methionine metabolism, glutathione metabolism and taurine/hypotaurine metabolism are among the most significantly different pathways (Supplementary Figure 5A). Notably, several molecules of the methionine cycle, transsulfuration and cysteine metabolism (L-methionine, S-adenosylmethionine - SAM, S-adenosylhomocysteine – SAH, L-cystathionine, hypotaurine) are significantly decreased in IDH1-mutant cells (Figure 3A, B). Oxidized glutathione (GSSG) is significantly higher in IDH1-mutant cells, while reduced glutathione (GSH) is significantly lower, as was also seen in end stage tumors by MEGA-PRESS MRS. A redox imbalance is also shown by the significantly lower GSH/GSSG ratio and the significantly lower hypotaurine/taurine ratio (Figure 3B). In line with these results and previous studies in human cell lines^30^, Seahorse metabolic assays also suggest that IDH1-mutant cells have significantly altered oxidative metabolism (Supplementary Figure 6). In addition, glutamate and glutamine, key molecules in cystine import and glutathione synthesis, are significantly enriched in IDH1-mutant cells, consistent with our *in vivo* MEGA-PRESS MRS results (Supplementary Figure 7A). Furthermore, our data shows that IDH1-mutant cells have significantly higher SAM/SAH ratio, indicating high methylation potential, and significantly lower cystathionine/SAH ratio (Figure 3B). These results suggest that IDH1-mutant cells have a metabolic state that favors the methionine cycle and methylation potential over transsulfuration, which can account for their increased vulnerability to cysteine deprivation.

**Figure 3.**
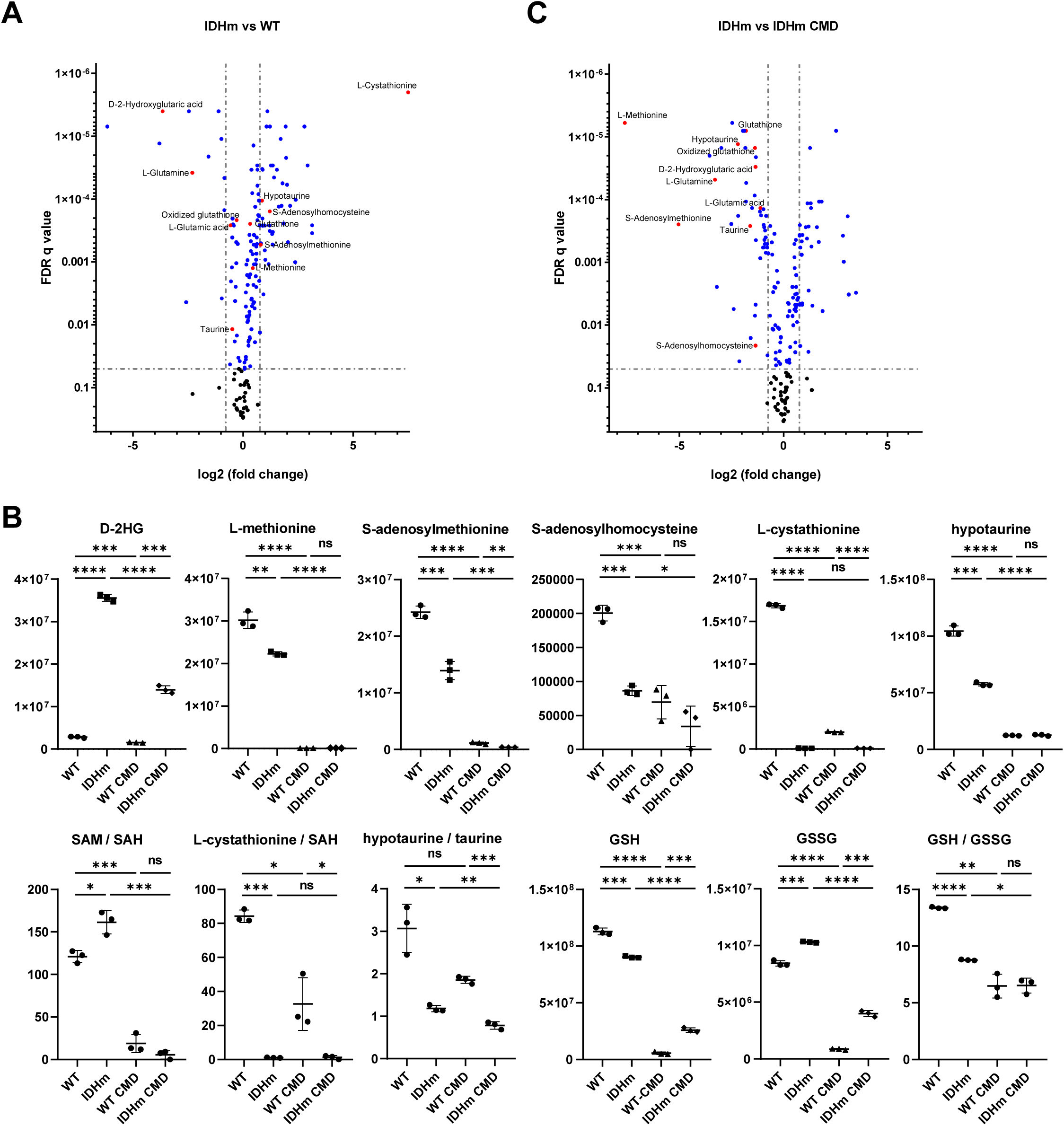
IDH1-mutant cells have an altered metabolic profile and are sensitive to cysteine-methionine deprivation. **A.** Volcano plot (blue and red: FDRq<0.05) of metabolomics analysis comparing IDH1-mutant cells (IDHm) to IDH1-wildtype (WT) cells after 17 hours in control conditions. **B.** Ratios and values of metabolites of the methionine cycle, transulfuration pathway and glutathione metabolism in untreated and CMD-treated IDHm and WT cells. Statistics assessed using t-test. Significance denoted by: *FDRq < 0.05, **FDRq < 0.01, ***FDRq < 0.001, ****FDRq<0.0001, ns: not significant (SAM: S-adenosylmethionine, SAH: S-adenosylhomocysteine, GSH: Glutathione, GSSG Oxidized glutathione). **C.** Volcano plot (blue and red: FDRq<0.05) comparing untreated IDHm cells to IDHm cells grown in CMD conditions for 17 hours.

To evaluate the metabolic effects of cysteine-methionine deprivation, we treated our mouse cell lines with CMD for 17 hours and then performed the same targeted metabolite profiling. Our results show that CMD significantly decreases metabolites involved in the methionine cycle, transsulfuration pathway and glutathione metabolism (Figure 3B, C, Supplementary Figures 5B, 7A), as well as the ratios SAM/SAH, GSH/GSSG and hypotaurine/taurine. These results indicate that CMD induces oxidative stress and disrupts transsulfuration in IDH1-wildtype cells, whereas IDH1-mutant cells show a pre-existing metabolic vulnerability due to altered cysteine and methionine metabolism, reduced transsulfuration, and increased oxidative stress, which is further exacerbated by CMD.

To characterize the metabolic effects of IDHmut-inhibitors, alone or in combination with CMD, we treated IDH1-mutant cells with 20mM IVO for 48 hours, CMD for 17 hours or combination of 20mM IVO for 48 hours with CMD for the last 17 hours of treatment (Figure 4, Supplementary Figures 7B, 8). In parallel, cells were treated with 1mM VOR for 48 hours (Supplementary Figure 9). Targeted metabolite profiling on more than 200 metabolites was performed across all conditions. CMD alone had the same effects as previously shown (Figure 3). Both IVO and VOR significantly decreased 2-HG, and altered many metabolites involved in cysteine/methionine metabolism (Figure 4A, Supplementary Figures 9A, B). Pathway analysis showed that IDHmut-inhibition significantly affects cysteine and methionine, glutathione, taurine and hypotaurine metabolism (Supplementary Figures 8, 9C). Specifically, IVO-treated cells have significantly decreased glutathione, as well as GSH/GSSG ratio, indicating higher oxidative stress. In addition, SAM and cystathionine are significantly decreased (Figure 4D). Similarly, in VOR-treated IDH1-mutant cells, SAM, cystathionine, glutathione and hypotaurine are significantly decreased (Supplementary Figure 9B). Combining CMD with IDHmut-inhibitors further exacerbated the effects on glutathione and the GSH/GSSG ratio (Figure 4B-D). Pathway analysis reveals that IDHmut-inhibition combined with CMD significantly affects glutathione, cysteine and methionine metabolism (Supplementary Figure 8). These metabolic changes provide a mechanistic correlation to our cell viability studies, supporting the finding that IDHmut-inhibition further sensitizes IDH1-mutant cells to CMD.

**Figure 4.**
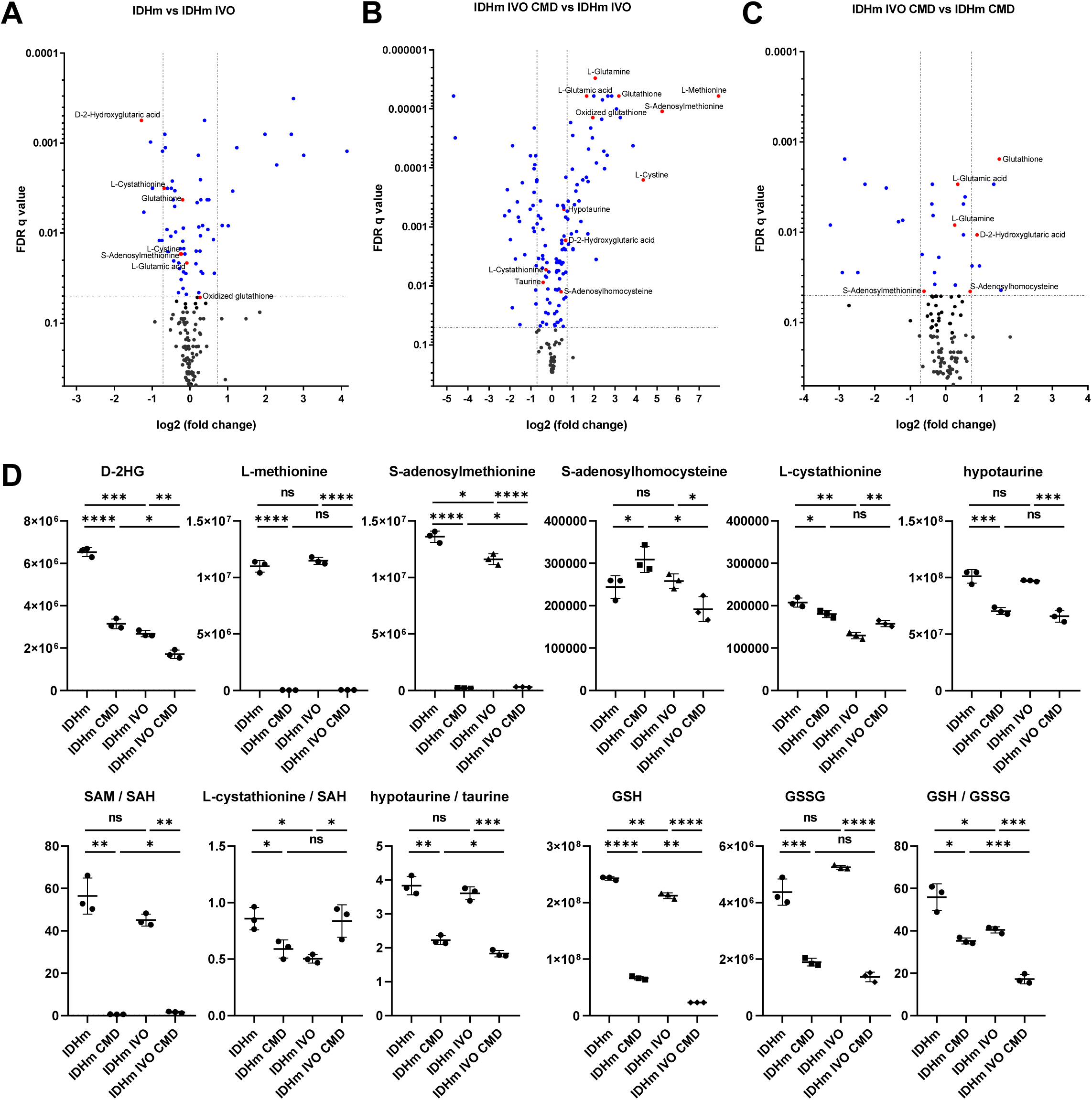
IDHmut-inhibitors enhance the sensitivity of IDH1-mutant glioma cells to cysteine-methionine deprivation. Volcano plots (blue and red: FDRq<0.05) of metabolomics results comparing **A.** untreated IDH1-mutant cells (IDHm) to treated with 20mM IVO for 48 hours (IDHm IVO), **B.** IDHm IVO to IDHm treated with 20mM IVO for 48 hours and CMD for 17 hours (IDHm IVO CMD) and **C.** IDHm IVO CMD to IDHm treated with CMD for 17 hours (IDHm CMD). **D.** Metabolite ratios and metabolites of the methionine cycle, transulfuration pathway and glutathione metabolism significantly altered by CMD and IVO treatments from experiments presented in A, B, C. Statistics assessed using t-test. Significance denoted by: *FDRq < 0.05, **FDRq < 0.01, ***FDRq < 0.001, ****FDRq<0.0001, ns: not significant (SAM: S-adenosylmethionine, SAH: S-adenosylhomocysteine, GSH: Glutathione, GSSG Oxidized glutathione).

### IDHmut-inhibition in combination with cysteine-methionine deprivation diet in vivo, prolongs survival of IDH1-mutant glioma-bearing mice

To assess whether IDH1-mutant glioma cells are more sensitive to CMD and ferroptosis *in vivo*, we tested the effects of RSL3 treatment in combination with a cysteine-deprived, methionine-restricted diet (CMD, 0.0% cysteine, 0.15% methionine) in mice with IDH1-mutant brain tumors. Notably, we treat cells with 0.0% cysteine and 0.0% methionine for CMD *in vitro*. For *in vivo* studies, methionine, as an essential amino acid, can be reduced but not completely removed from the diet. At 15dpi, mice were switched to either control diet or CMD and were maintained on these diets until end stage. Mice were treated with RSL3 or vehicle through convection-enhanced delivery (CED) for 7 days, starting at 30 dpi. Kaplan-Meier survival analysis showed a significant survival benefit for CMD or RSL3 treated mice over untreated mice. Notably, mice co-treated with both RSL3 and CMD survived significantly longer than control or CMD-treated mice (Figure 5A, B).

**Figure 5.**
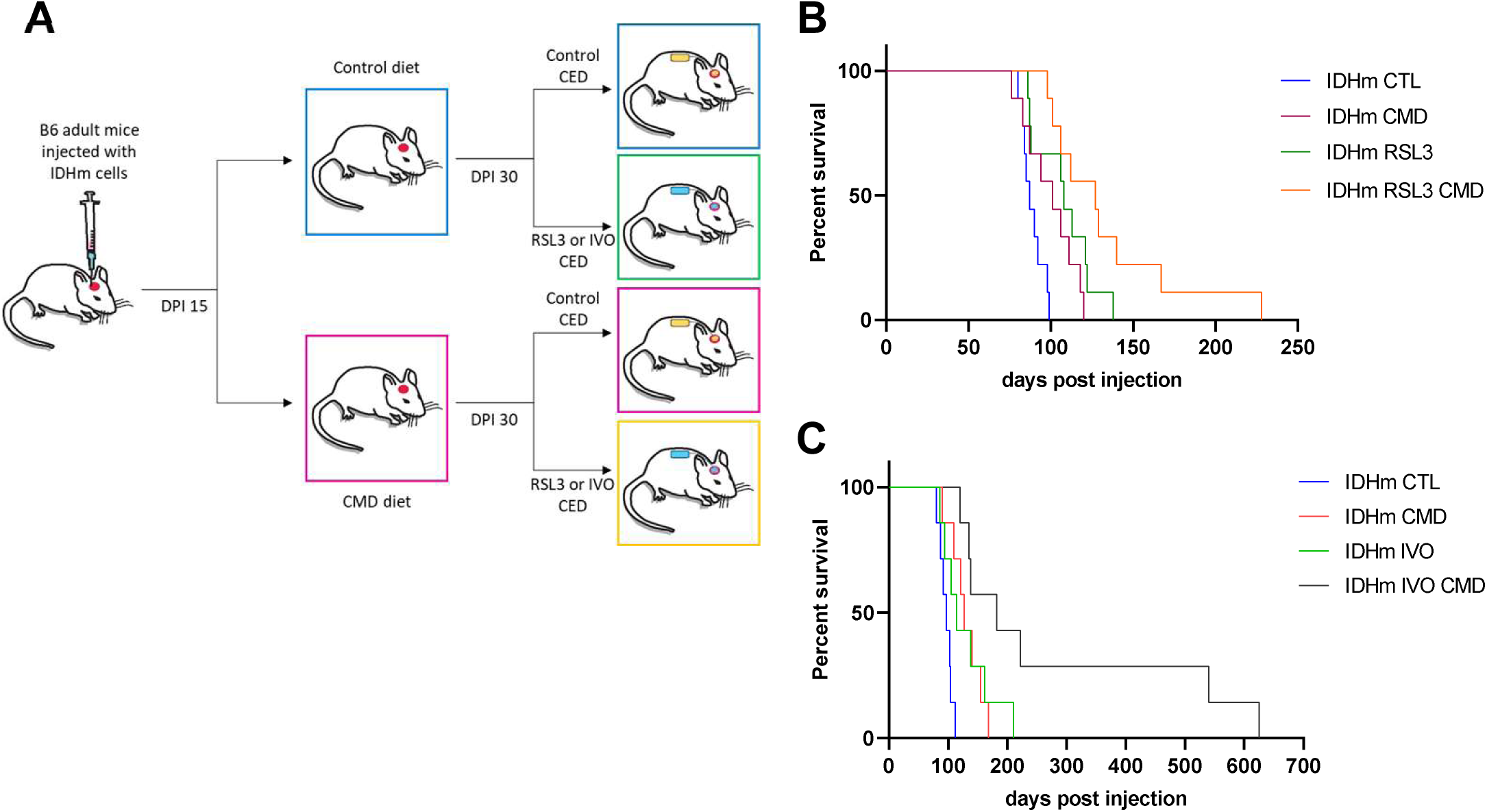
IDHmut-inhibition and cysteine-methionine deprivation prolong survival of IDH1-mutant tumor-bearing mice in vivo. **A.** Schematic representation of survival studies after intracranial injections of IDH1-mutant cells into adult mice. At 15 days post injection (dpi), mice were transferred to cysteine-deprived, methionine-restricted (CMD) or control diet. At 30dpi osmotic pumps were placed for intratumoral convection enhanced delivery (CED) for 7 days with 500nM RSL3 (RSL3 CED) or 0.01% DMSO (Control CED). In a second study, mice were treated for 7 days with 20mM IVO (IVO CED) or 0.01% DMSO. **B.** Survival curves after treatment with 500nM RSL3 and CMD in adult mice. Median survival: CTL 87dpi, CMD 101dpi, RSL3 108dpi, RSL3-CMD 127dpi. Log-Rank (Mantel-Cox) test: CTL vs RSL3 p=0.0050**, CTL vs CMD p=0.0280*, CMD vs RSL3 p=0.1563, RSL3 vs RSL3-CMD p=0.0743, CMD vs RSL3-CMD p=0.0104*, CTL vs RSL3-CMD p<0.0001****. **C.** Survival curves after treatment with 20mM IVO and CMD in adult mice. Median survival: CTL 97dpi, CMD 127dpi, IVO 114dpi, IVO-CMD 182dpi. Log-Rank (Mantel-Cox) test: CTL vs IVO p=0.0311*, CTL vs CMD p=0.0049**, CMD vs IVO p=0.9415, CMD vs IVO-CMD p=0.0417*, IVO vs IVO-CMD p=0.0449*, CTL vs IVO-CMD p=0.0001***.

To further explore the therapeutic potential of our findings, we tested the effects of IVO in combination with CMD diet in mice with IDH1-mutant brain tumors (Figure 5A, C). We selected ivosidenib (IVO) for these studies because it is highly selective for mutant IDH1^12,29^. Although clinical trials^12^ have shown that systemically administered vorasidenib (VOR) achieves superior brain penetrance compared to IVO, in our studies we employ CED, which bypasses the blood– brain barrier and permits direct intratumoral drug administration at substantially higher local concentrations than are achievable through systemic dosing. This approach allows us to maximize on-target inhibition within the tumor while minimizing systemic exposure and potential off-target or isoform-nonspecific effects. Using the same approach as in the CMD-RSL3 survival study above, at 15dpi mice were switched to either control or CMD diet and maintained on those diets for 7 months, until 230dpi (Figure 5A, C) when the remaining long-term survivors were switched from CMD to control diet. At 30dpi and for 7 days, mice were treated with CED of 20mM IVO or vehicle. Kaplan-Meier survival analysis showed a significant survival benefit for CMD or IVO treated mice over control mice. Notably, mice co-treated with both IVO and CMD survived significantly longer than control, IVO or CMD treated mice.

## Discussion

In this study we present a new mouse model of IDH1-mutant glioma, which successfully recapitulates the pathological characteristics of human low-grade IDH1-mutant gliomas, including diffuse infiltration, slow progression, and high levels of 2-HG. In addition, we identified significant metabolic alterations that were further exacerbated by cysteine-methionine deprivation and IDH1mut-inhibition. Previous studies have highlighted the limitations of establishing efficient IDH1-mutant glioma models *in vitro* and *in vivo*, that make studying the effects of the IDH1 mutation on metabolism and other cellular functions, as well as testing different therapeutic treatments, very challenging^31,32^. Here, comparing our IDH1-mutant model with an IDH1-wildtype model allowed us to identify key metabolic differences due to the presence of the IDH1 mutation. Our results reveal that IDH1-mutant cells are significantly more sensitive than IDH1-wildtype cells to cysteine deprivation and ferroptosis, and have significant alterations in glutathione, cysteine and methionine metabolism. Previous studies have focused on the epigenetic and metabolic alterations caused by 2-HG on glucose metabolism and the TCA cycle^33,34^, while studies on cysteine and methionine metabolism have shown that cystathionine, the transsulfuration intermediate, has low abundance in IDH1-mutant human glioma tissue and PDX models^13,35,36,37,38^. Furthermore, human IDH1-mutant glioma cells are sensitive to cysteine deprivation and can be rescued by the addition of cystathionine^35^. Our results demonstrate that cystathionine is significantly depleted in our mouse IDH1-mutant cells, recapitulating the human data and identifying deficiency in transsulfuration as a potential metabolic vulnerability for IDH1-mutant gliomas.

Our results reveal that, in addition to transsulfuration, IDH1-mutant gliomas have a dysfunction in the methionine cycle, as SAM is significantly lower in IDH1-mutant compared to IDH1-wildtype cells, and that is further exacerbated after treatments with CMD and IDHmut-inhibitors. It has previously been shown that elevated SAM levels directly promote transsulfuration^39,40,41^ and inhibit remethylation of homocysteine to methionine^17,42,43,44^. Compared to IDH1-wildtype, IDH1-mutant glioma cells have significantly lower levels of SAM and cystathionine, indicating disruption of the methionine cycle and transsulfuration pathway (Supplementary Figure 10). SAM levels decrease even further in CMD conditions, as well as after IDHmut-inhibition and further still after combination of both treatments. This coincides with further reduction of transsulfuration and diminished antioxidant defense, as demonstrated by the reduced ratios of GSH/GSSG and Hypotaurine/Taurine^45^. These effects render the IDH1-mutant cells more vulnerable to ferroptosis. Consistent with our results, a previous study showed that human IDH1-mutant glioma cells are sensitive to cysteine deprivation and depend on exogenous cysteine rather than transsulfuration for glutathione synthesis^35^. In our IDH1-mutant mouse model, exploiting this vulnerability in cysteine metabolism by combining IDHmut-inhibition and dietary CMD *in vivo*, significantly prolongs survival of tumor-bearing mice. Notably, while IDHmut-inhibitors significantly reduce 2-HG towards IDH1-wildtype levels, other metabolites remain altered or are further disturbed.

The synergy between CMD and IDHmut-inhibition may reflect the fact that pharmacologic inhibition of mutant IDH1 reverses some, but not all, metabolic consequences of the IDH1-mutant state. While IDHmut-inhibitors rapidly suppress 2-HG production, multiple studies have shown that broader metabolic, epigenetic, and redox-associated alterations persist or normalize only incompletely following treatment^46,47,48^. Concurrently, CMD acutely restricts cysteine availability and glutathione synthesis^36^, whereas restoration of redox homeostasis likely requires slower metabolic and transcriptional rewiring, including adjustments in transsulfuration and one-carbon metabolism, which can take several days^49,50^. Persistent or incompletely reversible alterations in sulfur metabolism or redox buffering may further enhance this vulnerability. Together, these findings suggest that combining dietary sulfur amino acid restriction with IDHmut-inhibition may exploit a transient metabolic liability in IDH1-mutant gliomas, providing a rationale for integrating CMD with IVO as a therapeutic strategy.

In conclusion, here we present a mouse model of IDH1-mutant glioma that can be used as a powerful tool to study the effects of both the IDH1 mutation and IDHmut-inhibitors *in vitro* and *in vivo*. Our results show that the IDH1-mutant cells have a disruption in cysteine and methionine metabolism and are sensitive to ferroptosis. IDHmut-inhibitors further exacerbate this vulnerability and, in combination with cysteine-methionine deprivation, induce oxidative stress and ferroptotic cell death of glioma cells *in vitro* and significantly prolong survival of tumor-bearing mice *in vivo*. Our study elucidates the benefits of combining IDHmut-inhibitors with a diet that targets specific metabolic vulnerabilities, in this case cysteine and methionine metabolism. Further studies are required to demonstrate efficiency in a clinical setting with the goal of improving therapeutic strategies for IDH1-mutant gliomas.

## Acknowledgments

This study was supported by the NIH/NCI (U54CA274504 and P01CA291697), the NIH/NINDS (R01NS103473), the Emerson Health Collective Cancer Research Fund and generous support from the Summer family. We thank Dr. Tak Wah Mak for the IDH1(R132H)^KI^ mouse. We thank Trine Giaever for her artistic help with the figures. Metabolomics mass spectrometric analysis was performed at the Weill Cornell Medicine Proteomics and Metabolomics Core Facility. The MRI imaging was carried out in the MR Facility of the Oncology Precision Therapeutics and Imaging Core (OPTIC) Shared Resource, supported by the Columbia University Medical Center NIH/NCI Cancer Center Support grant (P30CA013696). This study used the CCTI Flow Cytometry Core and the Molecular Pathology Shared Resource (MPSR) core, supported in part by the Office of the Director, NIH (S10OD020056). This work was supported by NIH Shared Instrumentation Grants (S10OD012351, S10OD021764 and S10OD032433), NIH/NCI Cancer Center Support grants (P30CA013696 and P30CA008748) and the Columbia University Digestive and Liver Disease Research Center grant (NIH 5P30DK132710).

## Author contributions

Conceptualization, A.Me. and P.C.; *In vivo* studies, A.Me., A.Ma. and N.H.; *In vitro* treatments A.Me., A.B., S.K., P.S.U.; qRT-PCR and MRI imaging A.Me. and A.Ma.; Immunohistochemistry A.Me., S.K., A.C.K.; Seahorse assays A.Me., A.B., T.T.T.N. and Q.G.; Metabolomics A.Me., A.B., S.L.; Flow cytometry analysis A.D.; MEGA-PRESS study J.G.; Interpretation of results A.Me., J.G., B.G., M.D.S., P.A.S., B.R.S., J.N.B. and P.C.; Writing – first draft A.Me. and P.C.; All authors provided critical feedback to shape the research, analysis and manuscript.

## Methods

### Mouse strains

Two mice (male and female) that harbor a knock-in (KI) R132H mutation of the mouse Idh1 gene in heterozygous condition^1^ were crossed to generate homozygous mice. These were then crossed with p53^fl/fl^; RiboTag^2^ mice and offspring were bred to Idh1(R132H)^KI^, p53^fl/fl^, and RiboTag homozygosity. These mice are viable and fertile as homozygotes. Since the Idh1(R132H) appears in heterozygous condition in human gliomas, for all necessary *in vivo* experiments we crossed Idh1(R132H)^KI^; p53^fl/fl^; RiboTag mice with +/+; p53^fl/fl^; RiboTag mice. The offspring have Idh1(R132H)^KI^ in heterozygous condition and maintain p53 and RiboTag as homozygotes^3,4^.

All experimental procedures involving mice were reviewed and approved by the Columbia University Institutional Animal Care and Use Committee (IACUC). The mice were monitored daily and euthanized when they exhibited signs that demonstrate that they are symptomatic from tumor burden including any of the following: decreased level of alertness and activity, seizures, periorbital hemorrhages, epistaxis (nose bleeds), impaired motor function, and/or impaired ability to feed secondary to decreased motor function, paresis or coma.

### Isolation of primary tumor cells and generation of cell lines

Murine glioma cell lines were generated from retrovirus-induced orthotopic murine glioma models. Neonatal (p4) Idh1(R132H)^KI^/+; p53^fl/fl^; RiboTag mice were anesthetized using hypothermia and orthotopically injected with 1μL of PDGFA or PDGFB–internal ribosomal entry site (IRES)–cyclization recombination (Cre) retrovirus with titer of 10^6^ cfu/μL and stereotactic coordinates relative to bregma: 1mm anterior, 1mm lateral, 1mm deep, aiming for subcortical white matter, using a Hamilton syringe at a flow rate of 0.3 μL/min. End-stage tumors were harvested and primary cells were isolated as follows: mice were sacrificed by cervical dislocation and decapitation, brains were acutely harvested and tumors were surgically resected. Tissue was minced and then dissociated in 10mL of 3% TrypLE (ThermoFisher Scientific, 12604013) in PBS, at 37°C under shaking for 10 minutes. Tissue was triturated using 1mL tips and subsequently filtered through a 70μm mesh. An equal volume of media was added to neutralize trypsin, and the suspension was centrifuged at 1200 rpm for 5 min. The supernatant was discarded, and the cell pellet was resuspended in media. The cell suspension was cultured in one well of a PLL-coated 6-well plate.

IDH1-wildtype tumor cells are cultured in basal media (BFP), containing DMEM (Gibco™ 11965-092) with 0.5% FBS (Gibco™ 16000044), 1% antibiotic-antimycotic (Gibco™ 15240-062), 1% N2 supplement (Gibco™ 17502-048), 10 ng/ml of recombinant human PDGF-AA (Peprotech 100-13A) and 10 ng/ml of recombinant human FGFb (Peprotech 100-18B-50UG).

IDH1-mutant tumor cells were isolated and cultured in RH media, an enriched form of BFP, based on media used for isolation of OPCs^5,6^, containing DMEM (Gibco™ 11965-092) with 0.25% FBS (Gibco™ 16000044), 1% antibiotic-antimycotic (Gibco™ 15240-062), 1.5% N2 supplement (Gibco™ 17502-048), 15 ng/ml of recombinant human PDGF-AA (Peprotech 100-13A), 15 ng/ml of recombinant human FGFb (Peprotech 100-18B-50UG), 2.5% Insulin-Transferrin-Selenium (Gibco™ 41400-045), 0.5% Sodium Pyruvate (SIGMA S8636), 0.5% GlutaMax™ (Gibco™ 35050-061), 0.5% B27 Supplement (Gibco™ 17504-044), 5 ng/ml biotin (SIGMA B4501). IDH1-mutant cells express high levels of 2-HG and grow better in RH medium (Supplementary Figure 11A, B).

A PDGFA-driven cell line made from one tumor with the IDH1R143H^KI^/+; p53^fl/fl^; RiboTag genetic background (IDHm or RHA), a PDGFB-expressing cell line made from one tumor with the same IDH1R143H^KI^/+; p53^fl/fl^; RiboTag background (IDHmB or RHB) and a PDGFB-expressing cell line from a tumor with +/+; p53^fl/fl^ background (WTB) were used for this study. A PDGFA-expressing +/+; p53^fl/fl^ cell line (WT, labeled MG3^7^ or APCL^8^ in previous studies) was also used. Similar results are observed with cell lines isolated from PDGFA-induced and PDGFB-induced primary tumors (IDHmB and WTB cell lines, Supplementary Figure 12).

### Orthotopic tumor implantation, diet allocation and convection enhanced delivery

Adherent cells were lifted using TrypLE (ThermoFisher Scientific, 12604013) and a cell suspension was made with a concentration of 7 × 10^4^ cells in 1μL. Mice were injected at 7-8 weeks old. Mice were anesthetized with Ketamine/Xylazine (100mg/kg and 10mg/kg, respectively) and assessed for lack of reflexes by toe pinch. Hair was shaved and scalp skin was incised. A burr hole was made with a 17-gauge needle 2mm lateral and 2mm anterior to bregma. Intracranial injection of 1μL of cell suspension was performed under stereotactic guidance, 2mm deep into the brain parenchyma aiming for subcortical white matter, using a Hamilton syringe at a flow rate of 0.3 μL/min. Tumor growth was assessed by MRI monitoring using a Bruker BioSpec 9.4 Tesla Small Animal MR Imager (CUIMC Oncology Precision Therapeutics and Imaging Core, OPTIC).

For *in vivo* studies with CMD diet, mice were transitioned to the diet fifteen days post tumor implantation. Special diets were created by LabTestDiet (W.F. Fisher and Sons): a control diet with 0.43% methionine and 0.33% cystine (w/w, catalog no. 5WVL) and a cystine-deprived, methionine-restricted diet with 0.15% methionine and 0.0% cystine (w/w, catalog no. 5WVM).

Similarly to what we have previously shown in the IDH-wildtype (WT) glioma model^7^, the diet was well tolerated throughout the duration of the survival studies, although notably CMD mice maintained lower weights than control mice. Investigators were not blinded to the diet allocation during experiments or outcome assessments.

After randomization to control or CMD diets, mice were further randomized into control or treatment groups using convection enhanced delivery. Pump implantation was performed at 30 days post tumor implantation. A total of 36 mice, 9 per group (5 female, 4 male), were included in the RSL3-CMD study and a total of 28 mice, 7 per group (4 female, 3 male), were included for the IVO-CMD study. Mini osmotic 7-day pumps (Alzet, model 1007D) were filled with 100μL of either 0.1% DMSO in PBS (control), 500nM RSL3 in PBS or 20mM ivosidenib in PBS. These pumps infuse 100 µL of fluid over 7 days. Mice were anesthetized with Ketamine/Xylazine (100mg/kg and 10mg/kg, respectively), hair was shaved and scalp skin incised. The pump was inserted subcutaneously on the back of the mouse, and a burr hole was made 1mm posterior to the tumor implantation site. Then, the CED catheter was inserted 3mm into the tumor and stabilized with instant adhesive (Loctite 454, Alzet #0008670). Pumps were explanted after 7 days of infusion (at 37dpi) and mice were monitored until they developed signs of tumor morbidity.

### Tissue processing

Mice were assessed daily for signs of tumor morbidity. At end stage, following cervical dislocation, the brain was surgically removed and processed according to the experiment. For histological analysis, the whole brain was placed in 4% PFA. After 24 hours, the brain was placed in PBS and sent to the Molecular Pathology Shared Resource (CUIMC MPSR) for paraffin embedding, sectioning and Hematoxylin-Eosin staining. For primary cell isolation, the largest anterior part of the tumor was used, and a small portion of the back was placed in 4% PFA to fix for histological analysis.

### Immunohistochemistry and antibodies

Paraffin-embedded 5μm tissue sections were processed for immunohistochemistry. Briefly, following deparaffinization with xylene and ethanol, sections were boiled in Sodium Citrate buffer, pH 6.0, for 10min at high pressure in a pressure cooker. After cooling down, sections were treated with 1% hydrogen peroxide to block endogenous peroxidases, then washed in PBS and incubated in blocking solution (0.5% Triton-X in PBS with 5% Horse Serum), at room temperature for 30-60 minutes. Following overnight (16-19 hours) incubation with primary antibody in blocking solution at 4°C, slides were washed in PBS and incubated with biotinylated secondary antibody for 1 hour and ABC reagent for 30-45 minutes according to the Vectastain Elite ABC kit instructions (Vector Laboratories). DAB (Dako) incubation was used to visualize staining, followed by counterstain with hematoxylin. Sections were then dehydrated and mounted with Permount. Primary antibodies for olig2 (Millipore AB9610, rabbit 1:200), Ki67 (Cell Signaling 9129, rabbit 1:200), GFAP (Dako Z0334, rabbit 1:500), NeuN (Millipore ABN78, rabbit 1:200), Iba1 (Cell Signaling 17198, rabbit 1:200), HA (Biolegend 901513, Covance MMS-101R clone 16B12, mouse IgG1, 1:500) were used.

### Cell culture and *in vitro* treatments

All cells were grown at 37°C with 5% CO2 in poly-L-lysine coated plates. All IDH1-wildtype cell lines were grown in BFP media. IDH1-mutant cells were grown in RH media. For all *in vitro* treatments, IDH1-mutant cell lines were transferred and grown into BFP media for 3 days before the experiment. Cysteine-methionine deprived (CMD) media was made from basal DMEM without cysteine, methionine, and glutamine (Thermo Fisher Scientific, 21013024) that was supplemented with L-glutamine to a final concentration of 4mM (as is the concentration in DMEM, Gibco™ 11965-092, used to make BFP). *In vitro* treatments of IDH1-mutant and IDH1-wildtype cells for 2 days with different concentrations (0-4mM) of glutamine, show that IDH1-mutant cells are significantly more dependent on glutamine (Supplementary Figure 11C). Cysteine deprived (CD) media was made from CMD media, supplemented with L-methionine to a final concentration of 30mg/L. All other components of the media were the same for control (BFP) and CMD media. Thus, for all experiments the only difference between CMD, CD and control conditions for each cell line was the concentration of cysteine and methionine in the media used. Murine glioma cells were plated in triplicate at a density of 7000 cells per well in a 96-well plate, then, 17 hours after plating, media was removed and treatment was added. We treated IDHm and WT cells *in vitro* with different concentrations of RSL3 (Stockwell lab), ivosidenib (MedChemExpress, AG-1205) or vorasidenib (MedChemExpress, AG-881) in control, CD or CMD media. Cell viability was assessed after treatment using a colorimetric reaction (MTS assay) with CellTiter 96 Aqueous One Solution reagent (Promega G3580). Absorbance at 490nm was detected with a BioRad Microplate Reader. Values were normalized to control or zero treatment conditions. Averages across 3 independent experiments are reported.

### 2-HG detection

To detect D-2-HG, 2×10^6^ IDH1-mutant cells were plated per 10cm plate coated with poly-L-lysine. After 48 hours, cells were lifted using TryPLE, and cell pellets were collected after centrifugation at 1200rpm for 5 minutes at room temperature. Cell pellets were frozen at -80°C and processed for 2-HG detection assay according to the instructions of D-2-Hydroxyglutarate Assay Kit (Abcam ab211070).

### Flow cytometric analysis of Lipid Peroxidation

IDH1-mutant and IDH1-wildtype cells were treated for 4 hours with either control or CMD media, followed by RSL3 treatment for 30 minutes at two different concentrations (50nM and 125nM). After treatment in 6-well plates, cells were lifted using TrypLE (ThermoFisher, 12604013). Cell pellets were resuspended in 1mL PBS with Bodipy-C11 (ThermoFisher, D3861) added to a final concentration of 2mM and incubated for 10min at 37°C. Cells were then washed once in PBS, centrifuged at 400 g for 5min, resuspended in flow buffer (PBS, 0.5% FBS, 1 μg/mL DAPI) and filtered in polystyrene flow tubes (Fisher Scientific #352008). Data were collected on a BD LSRIII Fortessa Cell Analyzer using Pacific Blue, FITC and PE filters as described^9^ and analyzed using FlowJo v10.

### Mitochondrial stress assay

The process using a Seahorse XFe24 analyzer is described in depth elsewhere^10^. A mitochondrial stress assay was performed based on Agilent Technologies manual. Murine glioma cell lines were seeded in XFe24 cell culture microplates (Agilent Technologies) at 18,000 cells per well in 250μL of BFP media. After 17 hours, media was aspirated and replaced with assay medium (Agilent Technologies). Mitochondrial stress tests were run in the following order: 2 μM oligomycin, 2 μM trifluoromethoxy carbonylcyanide phenylhydrazone (FCCP), and 0.5 μM rotenone/antimycin A.

### Metabolomics

To prepare samples for Metabolomics, 2×10^6^ cells were plated per 10cm plate coated with poly-L-lysine (3 plates per condition) and treated according to the experiment (BFP, CMD, BFP with IVO, BFP with IVO and CMD). Then, plates were washed 2 times with ice-cold PBS, aspirated, placed on dry ice and 1mL of 100% HPLC grade methanol was added to the dish. Cells were scraped and transferred with methanol into cold Eppendorf tubes. Samples were then centrifuged at 14,000 × g for 20min at 4°C. Pellets were used to extract protein using cell extraction buffer with protease and phosphatase inhibitors. A colorimetric Bradford assay was read at 740 nm for evaluation of total protein content. Supernatants were frozen at −80°C and sent in dry ice for mass spectrometric analysis at the Weill Cornell Medicine Proteomics and Metabolomics Core Facility. Samples were normalized by total metabolite counts and FDRq value was calculated in GraphPad Prism 10.0 with the two-stage step-up method of Benjamini, Krieger and Yekutieli^11^. Pathway analysis was performed using MetaboAnalyst 6.0 (https://www.metaboanalyst.ca/).

### Real time Quantitative PCR method

Primer sequences used for Slc7a11, Atf4 and Actin were as described previously^7^. Cells were treated into 6-well plates for 17 hours. Based on our *in vitro* viability studies, we used a prolonged treatment of 5nM RSL3 (close to IC50) to avoid cell death and detect possible treatment effects on mRNA transcripts. We treated IDH1-mutant and IDH1-wildtype cells with 5nM RSL3 with or without CMD for 17 hours. Cells were scraped into ice-cold PBS and cell pellets were collected after centrifugation at 14000rpm at 4°C and stored at -80°C. Total RNA was extracted from the cell pellets using the RNeasy Mini kit (QIAGEN, 74106). Up to 2.5 μg of RNA was used with the SuperScript Vilo cDNA synthesis kit (Thermo Fisher, 117540-50). cDNA was diluted to a concentration of 250 ng/μL and the RT-qPCR reactions were conducted with Thermo Scientific ABsolute Blue qPCR SYBR (ThermoFisher, AB4323A). Duplicate samples per condition were analyzed on an Applied Biosystems Quant Studio 3 qPCR instrument with all experiments being repeated 3 independent times. beta-Actin was used as reference and log fold change was calculated using the ddCT method comparing treatments to a control (BFP with 0.01% DMSO) sample.

### MR Spectroscopy

MRS MEGA-PRESS for 2-HG, Glx (glutamate and glutamine), glutathione and GABA was performed as described previously^12,13^.

## Notes

### Competing Interest Statement

B.R.S., P.C., J.N.B. are inventors on patents and patent applications involving ferroptosis. B.R.S. co-founded and serves as a consultant to ProJenX, Inc. and Exarta Therapeutics; holds equity in Sonata Therapeutics; serves as a consultant to Weatherwax Biotechnologies Corporation and Akin Gump Strauss Hauer & Feld LLP.

## References

1. Parsons DW, Jones S, Zhang X, Lin JC, Leary RJ, Angenendt P, Mankoo P, Carter H, Siu IM, Gallia GL, Olivi A, McLendon R, Rasheed BA, Keir S, Nikolskaya T, Nikolsky Y, Busam DA, Tekleab H, Diaz LA Jr, Hartigan J, Smith DR, Strausberg RL, Marie SK, Shinjo SM, Yan H, Riggins GJ, Bigner DD, Karchin R, Papadopoulos N, Parmigiani G, Vogelstein B, Velculescu VE, Kinzler KW. An integrated genomic analysis of human glioblastoma multiforme. Science. 2008 Sep 26;321(5897):1807–12. doi: 10.1126/science.1164382. PMID: 18772396; PMCID: PMC2820389.

2. Yan H, Parsons DW, Jin G, McLendon R, Rasheed BA, Yuan W, Kos I, Batinic-Haberle I, Jones S, Riggins GJ, Friedman H, Friedman A, Reardon D, Herndon J, Kinzler KW, Velculescu VE, Vogelstein B, Bigner DD. IDH1 and IDH2 mutations in gliomas. N Engl J Med. 2009 Feb 19;360(8):765–73. doi: 10.1056/NEJMoa0808710. PMID: 19228619; PMCID: PMC2820383.

3. Louis DN, Perry A, Wesseling P, Brat DJ, Cree IA, Figarella-Branger D, Hawkins C, Ng HK, Pfister SM, Reifenberger G, Soffietti R, von Deimling A, Ellison DW. The 2021 WHO Classification of Tumors of the Central Nervous System: a summary. Neuro Oncol. 2021 Aug 2;23(8):1231–1251. doi: 10.1093/neuonc/noab106. PMID: 34185076; PMCID: PMC8328013.

4. Dang L, White DW, Gross S, Bennett BD, Bittinger MA, Driggers EM, Fantin VR, Jang HG, Jin S, Keenan MC, Marks KM, Prins RM, Ward PS, Yen KE, Liau LM, Rabinowitz JD, Cantley LC, Thompson CB, Vander Heiden MG, Su SM. Cancer-associated IDH1 mutations produce 2-hydroxyglutarate. Nature. 2009 Dec 10;462(7274):739–44. doi: 10.1038/nature08617. PMID: 19935646; PMCID: PMC2818760.

5. Xu W, Yang H, Liu Y, Yang Y, Wang P, Kim SH, Ito S, Yang C, Wang P, Xiao MT, Liu LX, Jiang WQ, Liu J, Zhang JY, Wang B, Frye S, Zhang Y, Xu YH, Lei QY, Guan KL, Zhao SM, Xiong Y. Oncometabolite 2-hydroxyglutarate is a competitive inhibitor of α-ketoglutarate-dependent dioxygenases. Cancer Cell. 2011 Jan 18;19(1):17–30. doi: 10.1016/j.ccr.2010.12.014. PMID: 21251613; PMCID: PMC3229304.

6. Chowdhury R, Yeoh KK, Tian YM, Hillringhaus L, Bagg EA, Rose NR, Leung IK, Li XS, Woon EC, Yang M, McDonough MA, King ON, Clifton IJ, Klose RJ, Claridge TD, Ratcliffe PJ, Schofield CJ, Kawamura A. The oncometabolite 2-hydroxyglutarate inhibits histone lysine demethylases. EMBO Rep. 2011 May;12(5):463–9. doi: 10.1038/embor.2011.43. PMID: 21460794; PMCID: PMC3090014.

7. Losman JA, Kaelin WG Jr. What a difference a hydroxyl makes: mutant IDH, (R)-2-hydroxyglutarate, and cancer. Genes Dev. 2013 Apr 15;27(8):836–52. doi: 10.1101/gad.217406.113. PMID: 23630074; PMCID: PMC3650222.

8. Fu X, Chin RM, Vergnes L, Hwang H, Deng G, Xing Y, Pai MY, Li S, Ta L, Fazlollahi F, Chen C, Prins RM, Teitell MA, Nathanson DA, Lai A, Faull KF, Jiang M, Clarke SG, Cloughesy TF, Graeber TG, Braas D, Christofk HR, Jung ME, Reue K, Huang J. 2-Hydroxyglutarate Inhibits ATP Synthase and mTOR Signaling. Cell Metab. 2015 Sep 1;22(3):508–15. doi: 10.1016/j.cmet.2015.06.009. PMID: 26190651; PMCID: PMC4663076.

9. Ye D, Guan KL, Xiong Y. Metabolism, Activity, and Targeting of D- and L-2-Hydroxyglutarates. Trends Cancer. 2018 Feb;4(2):151–165. doi: 10.1016/j.trecan.2017.12.005. PMID: 29458964; PMCID: PMC5884165.

10. Mellinghoff IK, Ellingson BM, Touat M, Maher E, De La Fuente MI, Holdhoff M, Cote GM, Burris H, Janku F, Young RJ, Huang R, Jiang L, Choe S, Fan B, Yen K, Lu M, Bowden C, Steelman L, Pandya SS, Cloughesy TF, Wen PY. Ivosidenib in Isocitrate Dehydrogenase 1-Mutated Advanced Glioma. J Clin Oncol. 2020 Oct 10;38(29):3398–3406. doi: 10.1200/JCO.19.03327. PMID: 32530764; PMCID: PMC7527160.

11. Mellinghoff IK, Penas-Prado M, Peters KB, Burris HA 3rd, Maher EA, Janku F, Cote GM, de la Fuente MI, Clarke JL, Ellingson BM, Chun S, Young RJ, Liu H, Choe S, Lu M, Le K, Hassan I, Steelman L, Pandya SS, Cloughesy TF, Wen PY. Vorasidenib, a Dual Inhibitor of Mutant IDH1/2, in Recurrent or Progressive Glioma; Results of a First-in-Human Phase I Trial. Clin Cancer Res. 2021 Aug 15;27(16):4491–4499. doi: 10.1158/1078-0432.CCR-21-0611. PMID: 34078652; PMCID: PMC8364866.

12. Mellinghoff IK, Lu M, Wen PY, Taylor JW, Maher EA, Arrillaga-Romany I, Peters KB, Ellingson BM, Rosenblum MK, Chun S, Le K, Tassinari A, Choe S, Toubouti Y, Schoenfeld S, Pandya SS, Hassan I, Steelman L, Clarke JL, Cloughesy TF. Vorasidenib and ivosidenib in IDH1-mutant low-grade glioma: a randomized, perioperative phase 1 trial. Nat Med. 2023 Mar;29(3):615–622. doi: 10.1038/s41591-022-02141-2. Erratum in: Nat Med. 2024 Jan;30(1):302. doi: 10.1038/s41591-023-02473-7. PMID: 36823302; PMCID: PMC10313524.

13. Fack F, Tardito S, Hochart G, Oudin A, Zheng L, Fritah S, Golebiewska A, Nazarov PV, Bernard A, Hau AC, Keunen O, Leenders W, Lund-Johansen M, Stauber J, Gottlieb E, Bjerkvig R, Niclou SP. Altered metabolic landscape in IDH-mutant gliomas affects phospholipid, energy, and oxidative stress pathways. EMBO Mol Med. 2017 Dec;9(12):1681–1695. doi: 10.15252/emmm.201707729. PMID: 29054837; PMCID: PMC5709746.

14. Shi J, Sun B, Shi W, Zuo H, Cui D, Ni L, Chen J. Decreasing GSH and increasing ROS in chemosensitivity gliomas with IDH1 mutation. Tumour Biol. 2015 Feb;36(2):655–62. doi: 10.1007/s13277-014-2644-z. PMID: 25283382.

15. Mazor T, Chesnelong C, Pankov A, Jalbert LE, Hong C, Hayes J, Smirnov IV, Marshall R, Souza CF, Shen Y, Viswanath P, Noushmehr H, Ronen SM, Jones SJM, Marra MA, Cairncross JG, Perry A, Nelson SJ, Chang SM, Bollen AW, Molinaro AM, Bengtsson H, Olshen AB, Weiss S, Phillips JJ, Luchman HA, Costello JF. Clonal expansion and epigenetic reprogramming following deletion or amplification of mutant IDH1. Proc Natl Acad Sci U S A. 2017 Oct 3;114(40):10743–10748. doi: 10.1073/pnas.1708914114. PMID: 28916733; PMCID: PMC5635900.

16. Gilbert MR, Liu Y, Neltner J, Pu H, Morris A, Sunkara M, Pittman T, Kyprianou N, Horbinski C. Autophagy and oxidative stress in gliomas with IDH1 mutations. Acta Neuropathol. 2014 Feb;127(2):221–33. doi: 10.1007/s00401-013-1194-6. Epub 2013 Oct 23. PMID: 24150401; PMCID: PMC3946987.

17. Zhang HF, Klein Geltink RI, Parker SJ, Sorensen PH. Transsulfuration, minor player or crucial for cysteine homeostasis in cancer. Trends Cell Biol. 2022 Sep;32(9):800–814. doi: 10.1016/j.tcb.2022.02.009. PMID: 35365367; PMCID: PMC9378356.

18. Yu X, Long YC. Crosstalk between cystine and glutathione is critical for the regulation of amino acid signaling pathways and ferroptosis. Sci Rep. 2016 Jul 18;6:30033. doi: 10.1038/srep30033. PMID: 27425006; PMCID: PMC4948025.

19. Hayano M, Yang WS, Corn CK, Pagano NC, Stockwell BR. Loss of cysteinyl-tRNA synthetase (CARS) induces the transsulfuration pathway and inhibits ferroptosis induced by cystine deprivation. Cell Death Differ. 2016 Feb;23(2):270–8. doi: 10.1038/cdd.2015.93. PMID: 26184909; PMCID: PMC4716307.

20. Yang WS, Stockwell BR. Ferroptosis: Death by Lipid Peroxidation. Trends Cell Biol. 2016 Mar;26(3):165–176. doi: 10.1016/j.tcb.2015.10.014. PMID: 26653790; PMCID: PMC4764384.

21. Stockwell BR, Friedmann Angeli JP, Bayir H, Bush AI, Conrad M, Dixon SJ, Fulda S, Gascón S, Hatzios SK, Kagan VE, Noel K, Jiang X, Linkermann A, Murphy ME, Overholtzer M, Oyagi A, Pagnussat GC, Park J, Ran Q, Rosenfeld CS, Salnikow K, Tang D, Torti FM, Torti SV, Toyokuni S, Woerpel KA, Zhang DD. Ferroptosis: A Regulated Cell Death Nexus Linking Metabolism, Redox Biology, and Disease. Cell. 2017 Oct 5;171(2):273–285. doi: 10.1016/j.cell.2017.09.021. PMID: 28985560; PMCID: PMC5685180.

22. Hadian K, Stockwell BR. SnapShot: Ferroptosis. Cell. 2020 May 28;181(5):1188–1188.e1. doi: 10.1016/j.cell.2020.04.039. PMID: 32470402; PMCID: PMC8157339.

23. Upadhyayula PS, Higgins DM, Mela A, Banu M, Dovas A, Zandkarimi F, Patel P, Mahajan A, Humala N, Nguyen TTT, Chaudhary KR, Liao L, Argenziano M, Sudhakar T, Sperring CP, Shapiro BL, Ahmed ER, Kinslow C, Ye LF, Siegelin MD, Cheng S, Soni R, Bruce JN, Stockwell BR, Canoll P. Dietary restriction of cysteine and methionine sensitizes gliomas to ferroptosis and induces alterations in energetic metabolism. Nat Commun. 2023 Mar 2;14(1):1187. doi: 10.1038/s41467-023-36630-w. PMID: 36864031; PMCID: PMC9981683.

24. Gong T, Zhang X, Wei X, Yuan S, Saleh MG, Song Y, Edden RA, Wang G. GSH and GABA decreases in IDH1-mutated low-grade gliomas detected by HERMES spectral editing at 3 T *in vivo*. Neurochem Int. 2020 Dec;141:104889. doi: 10.1016/j.neuint.2020.104889. PMID: 33115694; PMCID: PMC7704685.

25. Pascuzzo R, Rudà R, Barker PB, Xiang J, Antelmi L, Gianeri R, Pellerino A, Mo F, Soffietti R, Bizzi A. Glutamate to GABA ratio is elevated in patients with IDH-mutant lower-grade gliomas and seizures. Neurooncol Adv. 2025 Aug 20;7(1):vdaf155. doi: 10.1093/noajnl/vdaf155. PMID: 40893414; PMCID: PMC12391666.

26. Wang N, Zeng GZ, Yin JL, Bian ZX. Artesunate activates the ATF4-CHOP-CHAC1 pathway and affects ferroptosis in Burkitt’s Lymphoma. Biochem Biophys Res Commun. 2019 Nov 12;519(3):533–539. doi: 10.1016/j.bbrc.2019.09.023. PMID: 31537387.

27. Fujii J, Homma T, Kobayashi S. Ferroptosis caused by cysteine insufficiency and oxidative insult. Free Radic Res. 2020 Dec;54(11-12):969–980. doi: 10.1080/10715762.2019.1666983. PMID: 31505959.

28. He F, Zhang P, Liu J, Wang R, Kaufman RJ, Yaden BC, Karin M. ATF4 suppresses hepatocarcinogenesis by inducing SLC7A11 (xCT) to block stress-related ferroptosis. J Hepatol. 2023 Aug;79(2):362–377. doi: 10.1016/j.jhep.2023.03.016. PMID: 36996941; PMCID: PMC11332364.

29. Barbato MI, Barone AK, Aungst SL, Miller CP, Ananthula S, Bi Y, Yang Y, Li X, Xiong Y, Fan J, Dorff SE, Zhao H, Zhou H, Pradhan S, Scepura B, Sinha AK, Stephenson M, Bhatnagar V, Saber H, Rahman NA, Tang S, Pazdur R, Kluetz PG, Larkins E, Drezner N. FDA Approval Summary: Vorasidenib for IDH-mutant Grade 2 Astrocytoma or Oligodendroglioma following surgery. Clin Cancer Res. 2025 Sep 5:10.1158/1078-0432.CCR-25-1333. doi: 10.1158/1078-0432.CCR-25-1333. PMID: 40911439; PMCID: PMC12416742.

30. Lu Y, Kwintkiewicz J, Liu Y, Tech K, Frady LN, Su YT, Bautista W, Moon SI, MacDonald J, Ewend MG, Gilbert MR, Yang C, Wu J. Chemosensitivity of IDH1-Mutated Gliomas Due to an Impairment in PARP1-Mediated DNA Repair. Cancer Res. 2017 Apr 1;77(7):1709–1718. doi: 10.1158/0008-5472.CAN-16-2773. PMID: 28202508; PMCID: PMC5380481.

31. Cohen AL, Holmen SL, Colman H. IDH1 and IDH2 mutations in gliomas. Curr Neurol Neurosci Rep. 2013 May;13(5):345. doi: 10.1007/s11910-013-0345-4. PMID: 23532369; PMCID: PMC4109985.

32. Tang LW, Mallela AN, Deng H, Richardson TE, Hervey-Jumper SL, McBrayer SK, Abdullah KG. Preclinical modeling of lower-grade gliomas. Front Oncol. 2023 Mar 27;13:1139383. doi: 10.3389/fonc.2023.1139383. PMID: 37051530; PMCID: PMC10083350.

33. Unruh D, Zewde M, Buss A, Drumm MR, Tran AN, Scholtens DM, Horbinski C. Methylation and transcription patterns are distinct in IDH mutant gliomas compared to other IDH mutant cancers. Sci Rep. 2019 Jun 20;9(1):8946. doi: 10.1038/s41598-019-45346-1. PMID: 31222125; PMCID: PMC6586617.

34. Han S, Liu Y, Cai SJ, Qian M, Ding J, Larion M, Gilbert MR, Yang C. IDH mutation in glioma: molecular mechanisms and potential therapeutic targets. Br J Cancer. 2020 May;122(11):1580–1589. doi: 10.1038/s41416-020-0814-x. PMID: 32291392; PMCID: PMC7250901.

35. Ruiz-Rodado V, Dowdy T, Lita A, Kramp T, Zhang M, Jung J, Dios-Esponera A, Zhang L, Herold-Mende CC, Camphausen K, Gilbert MR, Larion M. Cysteine is a limiting factor for glioma proliferation and survival. Mol Oncol. 2022 May;16(9):1777–1794. doi: 10.1002/1878-0261.13148. PMID: 34856072; PMCID: PMC9067152.

36. Cano-Galiano A, Oudin A, Fack F, Allega MF, Sumpton D, Martinez-Garcia E, Dittmar G, Hau AC, De Falco A, Herold-Mende C, Bjerkvig R, Meiser J, Tardito S, Niclou SP. Cystathionine-γ-lyase drives antioxidant defense in cysteine-restricted IDH1-mutant astrocytomas. Neurooncol Adv. 2021 Apr 9;3(1):vdab057. doi: 10.1093/noajnl/vdab057. PMID: 34250481; PMCID: PMC8262642.

37. Branzoli F, Pontoizeau C, Tchara L, Di Stefano AL, Kamoun A, Deelchand DK, Valabrègue R, Lehéricy S, Sanson M, Ottolenghi C, Marjańska M. Cystathionine as a marker for 1p/19q codeleted gliomas by *in vivo* magnetic resonance spectroscopy. Neuro Oncol. 2019 Jun 10;21(6):765–774. doi: 10.1093/neuonc/noz031. PMID: 30726924; PMCID: PMC6556848.

38. Branzoli F, Liserre R, Deelchand DK, Poliani PL, Bielle F, Nichelli L, Sanson M, Lehéricy S, Marjańska M. Neurochemical Differences between 1p/19q Codeleted and Noncodeleted IDH-mutant Gliomas by *in Vivo* MR Spectroscopy. Radiology. 2023 Sep;308(3):e223255. doi: 10.1148/radiol.223255. PMID: 37668523; PMCID: PMC10546286.

39. Pey AL, Majtan T, Sanchez-Ruiz JM, Kraus JP. Human cystathionine β-synthase (CBS) contains two classes of binding sites for S-adenosylmethionine (SAM): complex regulation of CBS activity and stability by SAM. Biochem J. 2013 Jan 1;449(1):109–21. doi: 10.1042/BJ20120731. PMID: 22985361.

40. McCorvie TJ, Kopec J, Hyung SJ, Fitzpatrick F, Feng X, Termine D, Strain-Damerell C, Vollmar M, Fleming J, Janz JM, Bulawa C, Yue WW. Inter-domain communication of human cystathionine β-synthase: structural basis of S-adenosyl-L-methionine activation. J Biol Chem. 2014 Dec 26;289(52):36018–30. doi: 10.1074/jbc.M114.610782. PMID: 25336647; PMCID: PMC4276868.

41. McCorvie TJ, Adamoski D, Machado RAC, Tang J, Bailey HJ, Ferreira DSM, Strain-Damerell C, Baslé A, Ambrosio ALB, Dias SMG, Yue WW. Architecture and regulation of filamentous human cystathionine beta-synthase. Nat Commun. 2024 Apr 4;15(1):2931. doi: 10.1038/s41467-024-46864-x. PMID: 38575566; PMCID: PMC10995199.

42. Finkelstein JD, Martin JJ. Homocysteine. Int J Biochem Cell Biol. 2000 Apr;32(4):385–9. doi: 10.1016/s1357-2725(99)00138-7. PMID: 10762063.

43. Hensley K, Denton TT. Alternative functions of the brain transsulfuration pathway represent an underappreciated aspect of brain redox biochemistry with significant potential for therapeutic engagement. Free Radic Biol Med. 2015 Jan;78:123–34. doi: 10.1016/j.freeradbiomed.2014.10.581. PMID: 25463282; PMCID: PMC4280296.

44. Sbodio JI, Snyder SH, Paul BD. Regulators of the transsulfuration pathway. Br J Pharmacol. 2019 Feb;176(4):583–593. doi: 10.1111/bph.14446. PMID: 30007014; PMCID: PMC6346075.

45. Karpowicz SJ. Kinetics of taurine biosynthesis metabolites with reactive oxygen species: Implications for antioxidant-based production of taurine. Biochim Biophys Acta Gen Subj. 2022 Jun;1866(6):130131. doi: 10.1016/j.bbagen.2022.130131. PMID: 35318954.

46. Rohle D, Popovici-Muller J, Palaskas N, Turcan S, Grommes C, Campos C, Tsoi J, Clark O, Oldrini B, Komisopoulou E, Kunii K, Pedraza A, Schalm S, Silverman L, Miller A, Wang F, Yang H, Chen Y, Kernytsky A, Rosenblum MK, Liu W, Biller SA, Su SM, Brennan CW, Chan TA, Graeber TG, Yen KE, Mellinghoff IK. An inhibitor of mutant IDH1 delays growth and promotes differentiation of glioma cells. Science. 2013 May 3;340(6132):626–30. doi: 10.1126/science.1236062. PMID: 23558169; PMCID: PMC3985613.

47. Turcan S, Makarov V, Taranda J, Wang Y, Fabius AWM, Wu W, Zheng Y, El-Amine N, Haddock S, Nanjangud G, LeKaye HC, Brennan C, Cross J, Huse JT, Kelleher NL, Osten P, Thompson CB, Chan TA. Mutant-IDH1-dependent chromatin state reprogramming, reversibility, and persistence. Nat Genet. 2018 Jan;50(1):62–72. doi: 10.1038/s41588-017-0001-z. PMID: 29180699; PMCID: PMC5769471.

48. Tateishi K, Wakimoto H, Iafrate AJ, Tanaka S, Loebel F, Lelic N, Wiederschain D, Bedel O, Deng G, Zhang B, He T, Shi X, Gerszten RE, Zhang Y, Yeh JJ, Curry WT, Zhao D, Sundaram S, Nigim F, Koerner MVA, Ho Q, Fisher DE, Roider EM, Kemeny LV, Samuels Y, Flaherty KT, Batchelor TT, Chi AS, Cahill DP. Extreme Vulnerability of IDH1 Mutant Cancers to NAD+ Depletion. Cancer Cell. 2015 Dec 14;28(6):773–784. doi: 10.1016/j.ccell.2015.11.006. PMID: 26678339; PMCID: PMC4684594.

49. Stone KP, Ghosh S, Kovalik JP, Orgeron M, Wanders D, Sims LC, Gettys TW. The acute transcriptional responses to dietary methionine restriction are triggered by inhibition of ternary complex formation and linked to Erk1/2, mTOR, and ATF4. Sci Rep. 2021 Feb 12;11(1):3765. doi: 10.1038/s41598-021-83380-0. PMID: 33580171; PMCID: PMC7880992.

50. Olsen T, Øvrebø B, Haj-Yasein N, Lee S, Svendsen K, Hjorth M, Bastani NE, Norheim F, Drevon CA, Refsum H, Vinknes KJ. Effects of dietary methionine and cysteine restriction on plasma biomarkers, serum fibroblast growth factor 21, and adipose tissue gene expression in women with overweight or obesity: a double-blind randomized controlled pilot study. J Transl Med. 2020 Mar 11;18(1):122. doi: 10.1186/s12967-020-02288-x. PMID: 32160926; PMCID: PMC7065370.

## References for Methods

1. Sasaki M, Knobbe CB, Itsumi M, Elia AJ, Harris IS, Chio II, Cairns RA, McCracken S, Wakeham A, Haight J, Ten AY, Snow B, Ueda T, Inoue S, Yamamoto K, Ko M, Rao A, Yen KE, Su SM, Mak TW. D-2-hydroxyglutarate produced by mutant IDH1 perturbs collagen maturation and basement membrane function. Genes Dev. 2012 Sep 15;26(18):2038–49. doi: 10.1101/gad.198200.112. PMID: 22925884; PMCID: PMC3444730.

2. Gonzalez C, Sims JS, Hornstein N, Mela A, Garcia F, Lei L, Gass DA, Amendolara B, Bruce JN, Canoll P, Sims PA. Ribosome profiling reveals a cell-type-specific translational landscape in brain tumors. J Neurosci. 2014 Aug 13;34(33):10924–36. doi: 10.1523/JNEUROSCI.0084-14.2014. PMID: 25122893; PMCID: PMC4131009.

3. Mela A, Dovas A, Mahajan A, Humala N, Nguyen T, Kanangat S, Upadhyayula P, Sprinzen L, Argenziano M, Huang D, Guo J, Casaccia P, Siegelin M, Stockwell B, Bruce J, Canoll P. EXTH-20. A mouse model reveals vulnerability of IDH1-mutated glioma to ferroptosis. Neuro Oncol. 2023 Nov 10;25(Suppl 5):v227–8. doi: 10.1093/neuonc/noad179.0873. PMCID: PMC10639641.

4. Mela A, Brand A, Mahajan A, Dovas A, Humala N, Kanangat S, Kleinstein A, Leskinen S, Nguyen TTT, Gao Q, Upadhyayula PS, Guo J, Gill BJA, Siegelin MD, Sims PA, Stockwell BR, Bruce JN, Canoll P. TMET-43. IDH1(R132H) inhibitors enhance the sensitivity of IDH1-mutant glioma to cysteine deprivation and ferroptosis. Neuro Oncol. 2025 Nov 11;27(Suppl 5):v438. doi: 10.1093/neuonc/noaf201.1734. PMCID: PMC12601626

5. Scaglione A, Patzig J, Liang J, Frawley R, Bok J, Mela A, Yattah C, Zhang J, Teo SX, Zhou T, Chen S, Bernstein E, Canoll P, Guccione E, Casaccia P. PRMT5-mediated regulation of developmental myelination. Nat Commun. 2018 Jul 19;9(1):2840. doi: 10.1038/s41467-018-04863-9. PMID: 30026560; PMCID: PMC6053423.

6. Sprinzen L, Garcia F, Mela A, Lei L, Upadhyayula P, Mahajan A, Humala N, Manier L, Caprioli R, Quiñones-Hinojosa A, Casaccia P, Canoll P. EZH2 Inhibition Sensitizes IDH1R132H-Mutant Gliomas to Histone Deacetylase Inhibitor. Cells. 2024 Jan 25;13(3):219. doi: 10.3390/cells13030219. PMID: 38334611; PMCID: PMC10854521.

7. Upadhyayula PS, Higgins DM, Mela A, Banu M, Dovas A, Zandkarimi F, Patel P, Mahajan A, Humala N, Nguyen TTT, Chaudhary KR, Liao L, Argenziano M, Sudhakar T, Sperring CP, Shapiro BL, Ahmed ER, Kinslow C, Ye LF, Siegelin MD, Cheng S, Soni R, Bruce JN, Stockwell BR, Canoll P. Dietary restriction of cysteine and methionine sensitizes gliomas to ferroptosis and induces alterations in energetic metabolism. Nat Commun. 2023 Mar 2;14(1):1187. doi: 10.1038/s41467-023-36630-w. PMID: 36864031; PMCID: PMC9981683.

8. Goldberg AR, Dovas A, Torres D, Pereira B, Viswanathan A, Das Sharma S, Mela A, Merricks EM, Megino-Luque C, McInvale JJ, Olabarria M, Shokooh LA, Zhao HT, Chen C, Kotidis C, Calvaresi P, Banu MA, Razavilar A, Sudhakar TD, Saxena A, Chokran C, Humala N, Mahajan A, Xu W, Metz JB, Bushong EA, Boassa D, Ellisman MH, Hillman EMC, Hargus G, Bravo-Cordero JJ, McKhann GM 2nd, Gill BJA, Rosenfeld SS, Schevon CA, Bruce JN, Sims PA, Peterka DS, Canoll P. Glioma-induced alterations in excitatory neurons are reversed by mTOR inhibition. Neuron. 2025 Jan 16. PMID: 39837324.

9. Banu MA, Dovas A, Argenziano MG, Zhao W, Sperring CP, Cuervo Grajal H, Liu Z, Higgins DM, Amini M, Pereira B, Ye LF, Mahajan A, Humala N, Furnari JL, Upadhyayula PS, Zandkarimi F, Nguyen TT, Teasley D, Wu PB, Hai L, Karan C, Dowdy T, Razavilar A, Siegelin MD, Kitajewski J, Larion M, Bruce JN, Stockwell BR, Sims PA, Canoll P. A cell state-specific metabolic vulnerability to GPX4-dependent ferroptosis in glioblastoma. EMBO J. 2024 Oct;43(20):4492–4521. doi: 10.1038/s44318-024-00176-4. PMID: 39192032; PMCID: PMC11480389.

10. Nguyen TTT, Zhang Y, Shang E, Shu C, Torrini C, Zhao J, Bianchetti E, Mela A, Humala N, Mahajan A, Harmanci AO, Lei Z, Maienschein-Cline M, Quinzii CM, Westhoff MA, Karpel-Massler G, Bruce JN, Canoll P, Siegelin MD. HDAC inhibitors elicit metabolic reprogramming by targeting super-enhancers in glioblastoma models. J Clin Invest. 2020 Jul 1;130(7):3699–3716. doi: 10.1172/JCI129049. PMID: 32315286; PMCID: PMC7324177.

11. Benjamini Y, Krieger AM, Yekutieli D. Adaptive linear step-up procedures that control the false discovery rate. Biometrika. 2006 Sep 1;93(3):491–507.

12. Ma DJ, Le HA, Ye Y, Laine AF, Lieberman JA, Rothman DL, Small SA, Guo J. MR spectroscopy frequency and phase correction using convolutional neural networks. Magn Reson Med. 2022 Apr;87(4):1700–1710. doi: 10.1002/mrm.29103. PMID: 34931715.

13. Guo J, Mela A, Qie Z, Yang Y, Mahajan A, Humala N, Canoll PD. Detecting IDH1 Mutations in Gliomas: Insights from J-Difference Editing MEGA-PRESS 1H-MRS. ISMRM & ISMRT annual meeting & exhibition, Singapore, May 2024, Abstract #0547. https://archive.ismrm.org/2024/0547.html.

