## Supplementary Figures 1-12 for "IDH-mutant inhibitors enhance the sensitivity of IDH1-mutant gliomas to cysteine-methionine deprivation and ferroptosis"

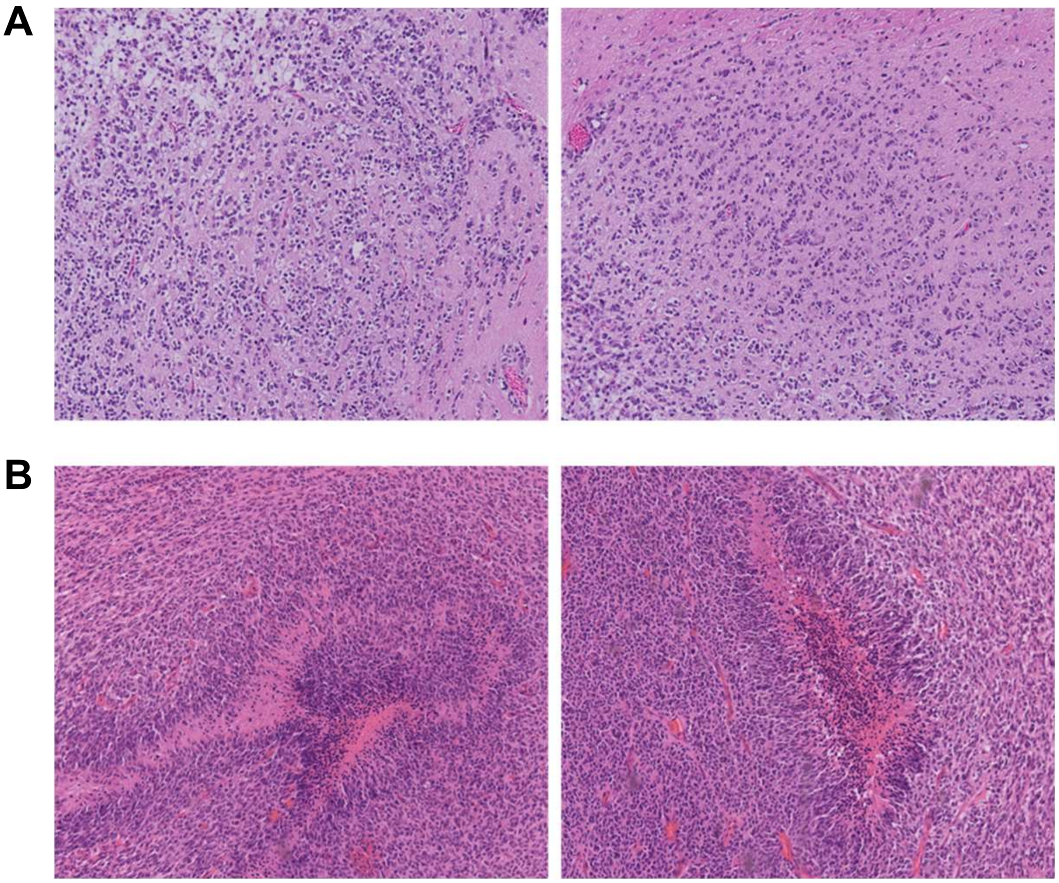

**Supplementary Figure 1. IDH1-mutant tumors have histopathological features of low-grade gliomas.**

Representative 10x H&E images of **A.** IDH1-mutant (IDH1-PDGFA) end stage mouse gliomas showing diffuse infiltration or **B.** IDH1-wildtype (WT-PDGFA) end stage mouse gliomas showing palisading necrosis and hemorrhage.

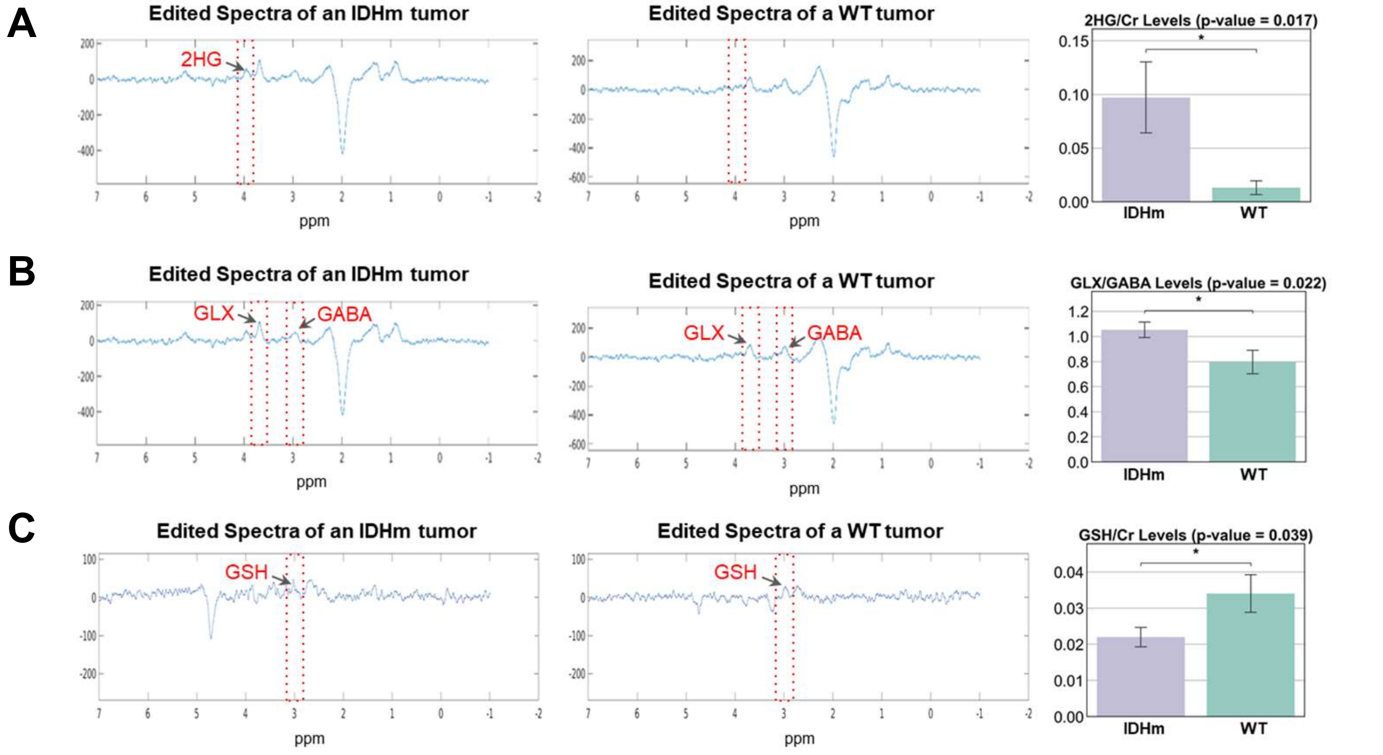


**Supplementary Figure 2. IDH1-mutant mouse tumors express 2-HG, have low glutathione and high GLX/GABA *in vivo*.**

MEGA-PRESS results after *in vivo* imaging of IDH1-mutant (IDHm) and IDH1-wildtype (WT) mouse end stage tumors, detecting **A.** 2-HG (p=0.017, IDHm n=12, WT n=6), **B.** GLX and GABA (p=0.022, IDHm n=10, WT n=11), or **C.** GSH (p=0.039, IDHm n=12, WT n=8). Data is presented as mean ± SD. Statistics assessed using t-test. Significance denoted by: *p < 0.05.

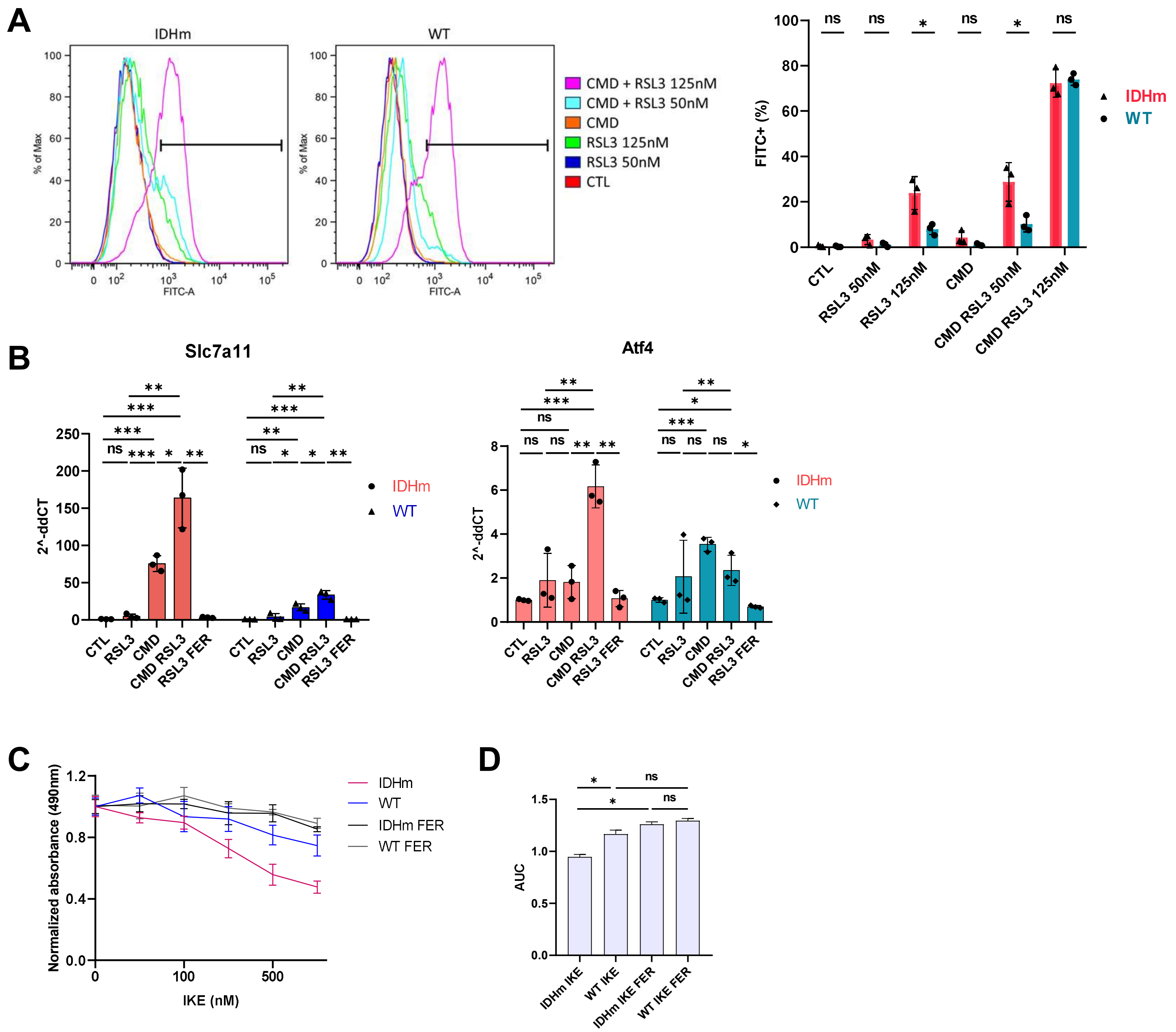


**Supplementary Figure 3. IDH1-mutant glioma cells are sensitive to RSL3, IKE and CMD-induced ferroptosis.**

**A.** Flow cytometry results after Bodipy C-11. IDH1-mutant (IDHm) and IDH1-wildtype (WT) cells were pre-treated with control (BFP) or CMD media for 4 hours, then with RSL3 for 30 minutes, followed by Bodipy-C11 staining and flow cytometry. Data presented as mean $\pm$ SD. Statistics assessed using t-test. Significance denoted by: $*p < 0.05$, ns: not significant. **B.** qRT-PCR results for Slc7a11 and Atf4 transcripts. IDHm and WT cells were treated for 17 hours in control or CMD media with 0.01% DMSO (BFP or CMD) or 5nM RSL3 (RSL3 or CMD RSL3) or 5nM RSL3 and 2µM Ferrostatin-1 (RSL3 FER). Data plotted as 2^-ddCT ± SEM, t-test significance denoted by: $*p < 0.05$, $**p < 0.01$, $***p < 0.001$, ns: not significant. **C.** Dose response curves after 24-hour treatment with IKE with or without 2µM Ferrostatin. Data plotted as normalized mean $\pm$ SD. IKE concentrations tested 0nM, 50nM, 100nM, 250nM, 500nM, 1000nM. **D.** Area under the curve (AUC) plotted from three independent experiments as presented in C. t-test significance denoted by: $*p < 0.05$, ns: not significant.

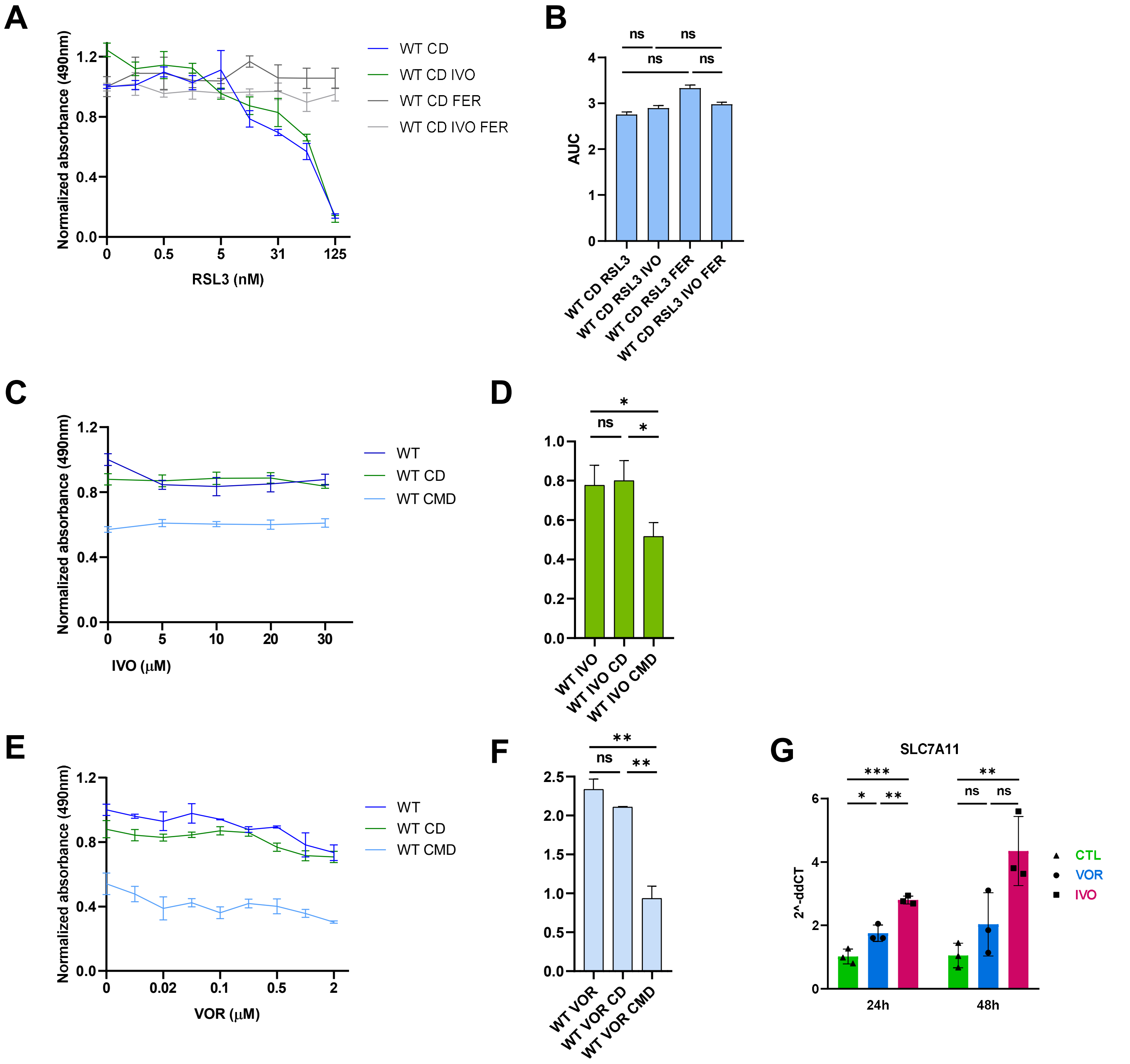


**Supplementary Figure 4. IDHmut-inhibitors do not significantly affect IDH1-wildtype cells and increase Slc7a11 in IDH1-mutant cells.**

**A.** Dose response curves of IDH1-wildtype (WT) cells after treatments with RSL3 (0, 0.1, 0.5, 1, 5, 10, 31, 62, 125 nM) in CD conditions with or without 20μM IVO or Ferrostatin-1 (FER) for 17 hours (IC50 WT CD 132.4, WT CD IVO 129.3). **B.** Area under the curve (AUC) plotted from three independent experiments as presented in A. **C.** Dose response curves after 17-hour treatment of IDH1-wildtype cells with IVO (0, 5, 10, 20, 30 μM) in control, CD or CMD conditions. **D.** Area under the curve (AUC) plotted from three independent experiments as presented in C. **E.** Dose response curves after 17-hour treatment of IDH1-wildtype cells with VOR (0, 0.005, 0.02, 0.05, 0.1, 0.2, 0.5, 1, 2 μM) in control, CD or CMD conditions. **F.** Area under the curve (AUC) plotted from three independent experiments as presented in E. Data plotted as normalized mean $\pm$ SD, t-test significance denoted by: *$p < 0.05$, **$p < 0.01$, ***$p < 0.001$, ****$p<0.0001$, ns: not significant. **G.** qRT-PCR results for Slc7a11 transcripts in IDH1-mutant cells treated for 24 or 48 hours with 0.01% DMSO (CTL) or 20μM IVO or 0.1μM VOR. Data plotted as 2^-ddCT ± SEM, t-test significance denoted by: *$p < 0.05$, **$p < 0.01$, ***$p < 0.001$. ns: not significant.

**A**

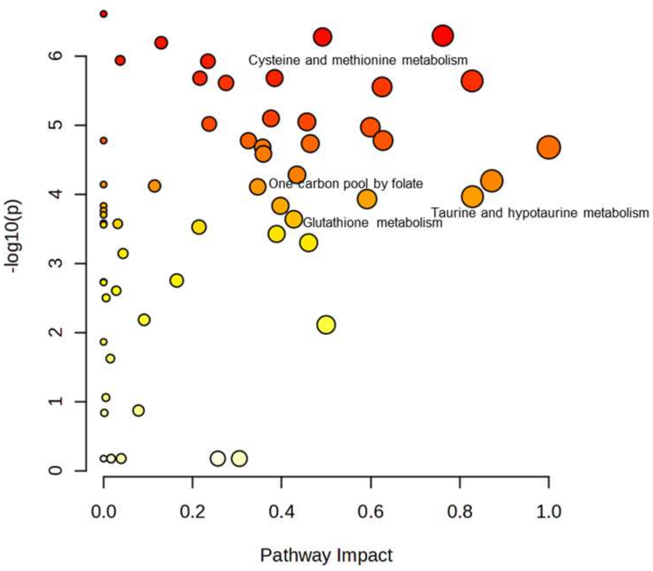


| Pathway Name | Match Status | p | -log(p) | Holm p | FDR | Impact |
|---|---|---|---|---|---|---|
| Ether lipid metabolism | 1/20 | 2.47E-07 | 6.6075 | 1.46E-05 | 9.50E-06 | 0 |
| Ascorbate and aldarate metabolism | 4/9 | 5.09E-07 | 6.2932 | 2.95E-05 | 9.50E-06 | 0.76191 |
| Arginine and proline metabolism | 13/36 | 5.31E-07 | 6.2746 | 3.03E-05 | 9.50E-06 | 0.49185 |
| Inositol phosphate metabolism | 5/30 | 6.44E-07 | 6.1909 | 3.61E-05 | 9.50E-06 | 0.12939 |
| Galactose metabolism | 4/27 | 1.16E-06 | 5.9365 | 6.37E-05 | 1.17E-05 | 0.03727 |
| Glycerophospholipid metabolism | 8/36 | 1.19E-06 | 5.9232 | 6.44E-05 | 1.17E-05 | 0.2343 |
| Cysteine and methionine metabolism | 7/33 | 2.09E-06 | 5.68 | 1.11E-04 | 1.45E-05 | 0.38414 |
| beta-Alanine metabolism | 7/21 | 2.10E-06 | 5.6783 | 1.11E-04 | 1.45E-05 | 0.21642 |
| Alanine, aspartate and glutamate metabolism | 16/28 | 2.29E-06 | 5.6394 | 1.17E-04 | 1.45E-05 | 0.82773 |
| Glyoxylate and dicarboxylate metabolism | 9/32 | 2.46E-06 | 5.6096 | 1.23E-04 | 1.45E-05 | 0.275 |
| One carbon pool by folate | 10/26 | 7.82E-05 | 4.1069 | 0.00258 | 1.71E-04 | 0.34627 |
| Taurine and hypotaurine metabolism | 2/8 | 1.08E-04 | 3.9649 | 0.00347 | 2.28E-04 | 0.82857 |
| Glutathione metabolism | 9/28 | 2.30E-04 | 3.6389 | 0.005971 | 3.99E-04 | 0.4272 |

**B**

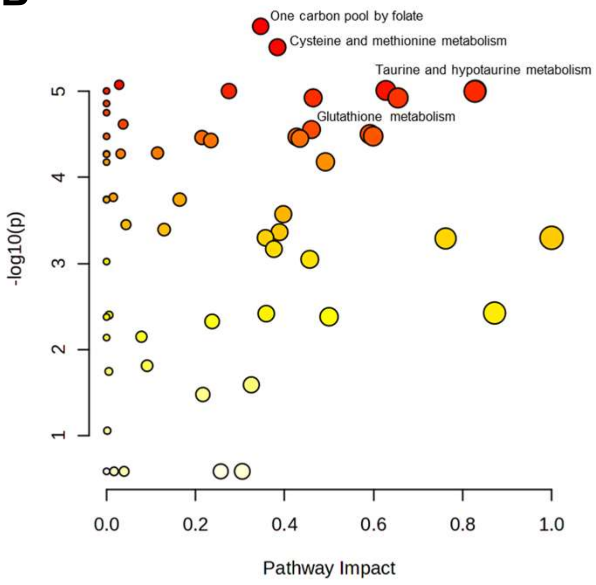


| Pathway Name | Match Status | p | -log(p) | Holm p | FDR | Impact |
|---|---|---|---|---|---|---|
| One carbon pool by folate | 10/26 | 1.75E-06 | 5.7558 | 1.04E-04 | 7.03E-05 | 0.34627 |
| Cysteine and methionine metabolism | 7/33 | 3.07E-06 | 5.5125 | 1.78E-04 | 7.03E-05 | 0.38414 |
| Valine, leucine and isoleucine degradation | 4/40 | 8.39E-06 | 5.0763 | 4.78E-04 | 7.03E-05 | 0.02836 |
| Arginine biosynthesis | 11/14 | 9.76E-06 | 5.0107 | 5.46E-04 | 7.03E-05 | 0.62766 |
| Taurine and hypotaurine metabolism | 2/8 | 9.90E-06 | 5.0045 | 5.46E-04 | 7.03E-05 | 0.82857 |
| Nitrogen metabolism | 2/6 | 9.91E-06 | 5.0041 | 5.46E-04 | 7.03E-05 | 0 |
| Glyoxylate and dicarboxylate metabolism | 9/32 | 9.91E-06 | 5.0037 | 5.46E-04 | 7.03E-05 | 0.275 |
| Alanine, aspartate and glutamate metabolism | 16/28 | 1.00E-05 | 4.9994 | 5.46E-04 | 7.03E-05 | 0.82773 |
| Purine metabolism | 20/70 | 1.19E-05 | 4.9245 | 6.07E-04 | 7.03E-05 | 0.46423 |
| Pyrimidine metabolism | 21/39 | 1.19E-05 | 4.924 | 6.07E-04 | 7.03E-05 | 0.65502 |
| Glutathione metabolism | 9/28 | 3.38E-05 | 4.4714 | 0.001463 | 1.04E-04 | 0.4272 |

**Supplementary Figure 5. Cysteine-methionine metabolism is significantly altered in IDH1-mutant cells.** MetaboAnalyst 6.0 (KEGG) metabolite pathway analysis comparing **A.** untreated IDH1-mutant and IDH1-wildtype cells, or **B.** untreated and CMD-treated IDH1-mutant cells. Top significantly affected pathways are shown with highlighted glutathione, cysteine and methionine metabolism-related pathways.

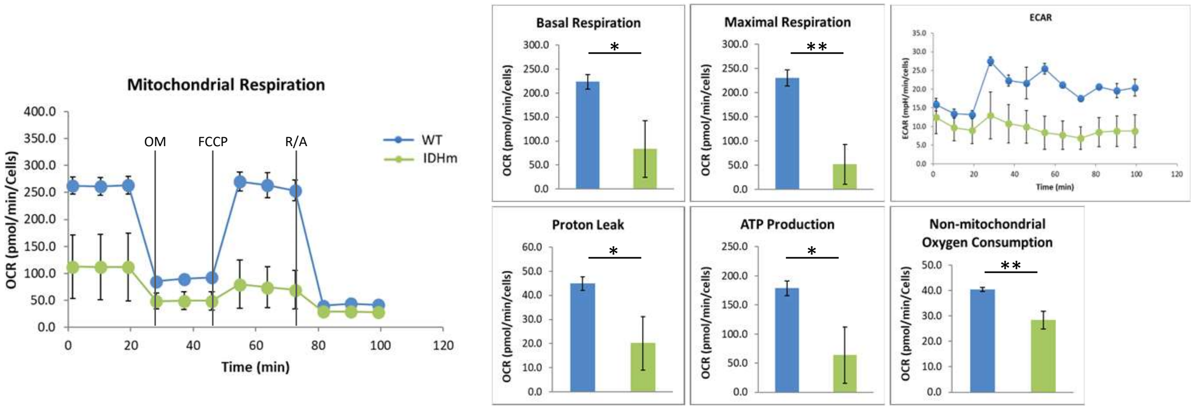


**Supplementary Figure 6. IDH1-mutant cells are sensitive to oxidative stress.**
Seahorse Mitochondrial stress test of IDH1-wildtype (WT: blue) and IDH1-mutant (IDHm: green) cells in BFP media (n = 4 per group). OM: oligomycin, FCCP: Carbonyl cyanide-4 (trifluoromethoxy) phenylhydrazone, R/A: rotenone and antimycin A. Basal respiration, maximal respiration, ATP production, non-mitochondrial oxygen consumption and proton leak values were calculated and normalized. Extracellular acidification rate (ECAR) is also presented. Data is presented as mean ± SD. Statistics assessed using t-test. Significance denoted by: $*p < 0.05$, $**p < 0.01$.

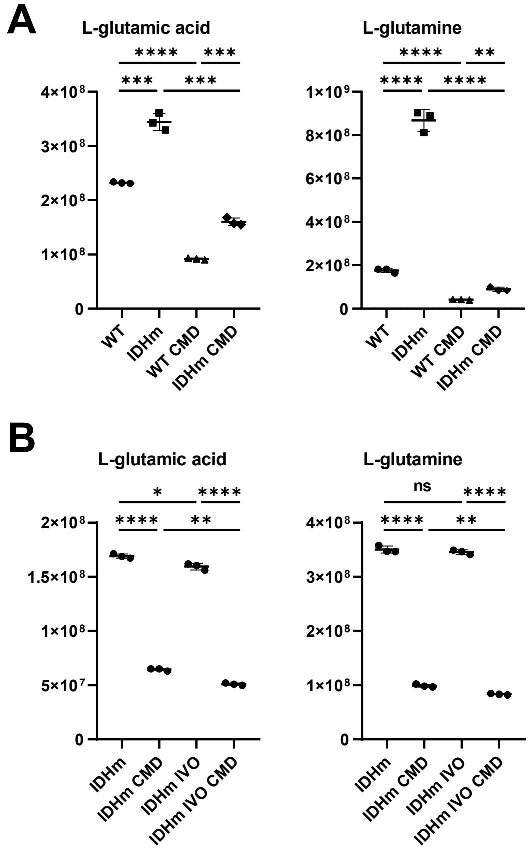


**Supplementary Figure 7. Glutathione metabolism-related metabolites are affected by CMD and ivosidenib treatments.**

Metabolomics results for L-glutamic acid and L-glutamine comparing **A.** untreated and CMD-treated IDH1-mutant (IDHm) and IDH1-wildtype (WT) cells, **B.** untreated and CMD-treated or IVO-treated or combo-treated IDH1-mutant cells (20μM IVO for 48 hours, CMD for 17 hours). Statistics assessed using t-test. Significance denoted by: *FDRq < 0.05, **FDRq < 0.01, ***FDRq < 0.001, ****FDRq<0.0001, ns: not significant.

A

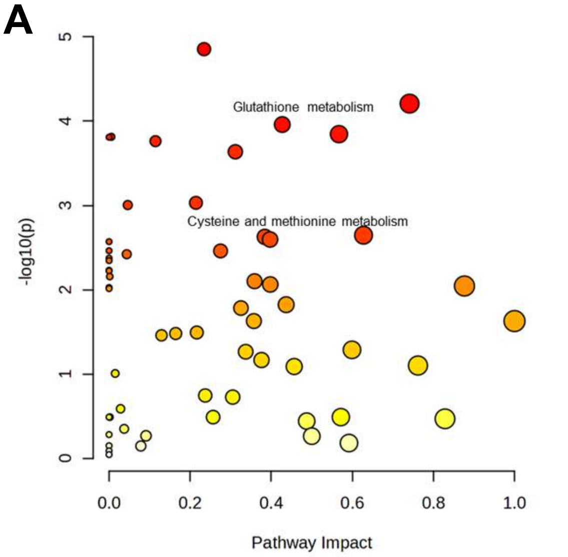


| Pathway Name | Match Status | p | -log(p) | Holm p | FDR | Impact |
|---|---|---|---|---|---|---|
| Glycerophospholipid metabolism | 8/36 | 1.40E-05 | 4.8533 | 8.13E-04 | 8.13E-04 | 0.2343 |
| Alanine, aspartate and glutamate metabolism | 15/28 | 6.20E-05 | 4.2079 | 0.003532 | 0.001432 | 0.74119 |
| Glutathione metabolism | 9/28 | 1.10E-04 | 3.9595 | 0.006147 | 0.001432 | 0.4272 |
| Nicotinate and nicotinamide metabolism | 6/15 | 1.43E-04 | 3.8458 | 0.007844 | 0.001432 | 0.56711 |
| Sphingolipid metabolism | 2/32 | 1.54E-04 | 3.8139 | 0.008289 | 0.001432 | 0.00563 |
| D-Amino acid metabolism | 1/15 | 1.55E-04 | 3.8094 | 0.008289 | 0.001432 | 0 |
| Lysine degradation | 7/30 | 1.73E-04 | 3.7625 | 0.008984 | 0.001432 | 0.11451 |
| Histidine metabolism | 6/16 | 2.31E-04 | 3.6367 | 0.011773 | 0.001674 | 0.31147 |
| Pantothenate and CoA biosynthesis | 6/20 | 9.30E-04 | 3.0314 | 0.046517 | 0.005731 | 0.21429 |
| Propanoate metabolism | 3/22 | 9.88E-04 | 3.0052 | 0.048415 | 0.005731 | 0.04616 |
| Cysteine and methionine metabolism | 8/33 | 0.002356 | 2.6279 | 0.11072 | 0.011127 | 0.38414 |

B

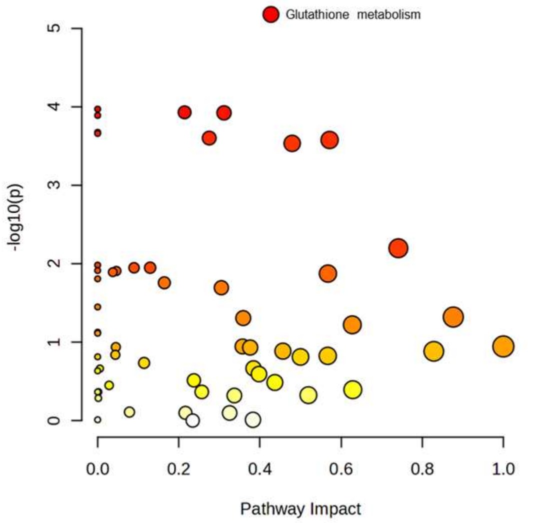


| Pathway Name | Match Status | p | -log(p) | Holm p | FDR | Impact |
|---|---|---|---|---|---|---|
| Glutathione metabolism | 9/28 | 6.68E-06 | 5.1753 | 3.87E-04 | 3.87E-04 | 0.4272 |
| Butanoate metabolism | 8/15 | 1.07E-04 | 3.9692 | 0.006119 | 0.001499 | 0 |
| Pantothenate and CoA biosynthesis | 6/20 | 1.18E-04 | 3.9283 | 0.006605 | 0.001499 | 0.21429 |
| Histidine metabolism | 6/16 | 1.20E-04 | 3.9214 | 0.006605 | 0.001499 | 0.31147 |
| Porphyrin metabolism | 2/31 | 1.29E-04 | 3.8887 | 0.006978 | 0.001499 | 0 |
| Thiamine metabolism | 1/7 | 2.12E-04 | 3.6733 | 0.011246 | 0.001701 | 0 |
| Nitrogen metabolism | 2/6 | 2.19E-04 | 3.6593 | 0.011395 | 0.001701 | 0 |
| Glyoxylate and dicarboxylate metabolism | 10/32 | 2.51E-04 | 3.6002 | 0.012804 | 0.001701 | 0.275 |
| Pyrimidine metabolism | 21/39 | 2.66E-04 | 3.5759 | 0.013275 | 0.001701 | 0.57171 |
| Purine metabolism | 20/70 | 2.93E-04 | 3.5326 | 0.014373 | 0.001701 | 0.48749 |

C

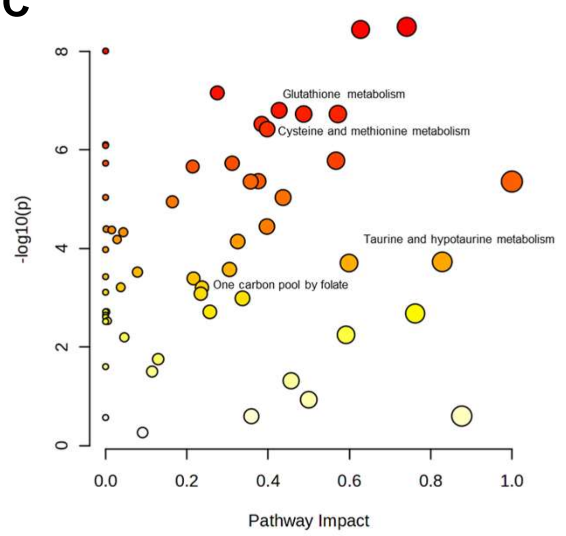


| Pathway Name | Match Status | p | -log(p) | Holm p | FDR | Impact |
|---|---|---|---|---|---|---|
| Alanine, aspartate and glutamate metabolism | 15/28 | 3.16E-09 | 8.5 | 1.83E-07 | 1.04E-07 | 0.74119 |
| Arginine biosynthesis | 11/14 | 3.58E-09 | 8.4462 | 2.04E-07 | 1.04E-07 | 0.62766 |
| Nitrogen metabolism | 2/6 | 9.80E-09 | 8.0086 | 5.49E-07 | 1.90E-07 | 0 |
| Glyoxylate and dicarboxylate metabolism | 10/32 | 6.93E-08 | 7.1593 | 3.81E-06 | 1.00E-06 | 0.275 |
| Glutathione metabolism | 9/28 | 1.57E-07 | 6.8042 | 8.48E-06 | 1.55E-06 | 0.4272 |
| Purine metabolism | 20/70 | 1.86E-07 | 6.7293 | 9.88E-06 | 1.55E-06 | 0.48749 |
| Pyrimidine metabolism | 21/39 | 1.87E-07 | 6.7288 | 9.88E-06 | 1.55E-06 | 0.57171 |
| Cysteine and methionine metabolism | 8/33 | 2.98E-07 | 6.5259 | 1.52E-05 | 2.16E-06 | 0.38414 |
| Arginine and proline metabolism | 11/36 | 3.78E-07 | 6.4227 | 1.89E-05 | 2.44E-06 | 0.39767 |
| Butanoate metabolism | 4/15 | 7.79E-07 | 6.1083 | 3.82E-05 | 4.29E-06 | 0 |
| Taurine and hypotaurine metabolism | 2/8 | 1.87E-04 | 3.7284 | 0.005606 | 3.74E-04 | 0.82857 |
| One carbon pool by folate | 10/26 | 0.001029 | 2.9877 | 0.020574 | 0.00153 | 0.33697 |

**Supplementary Figure 8. CMD and ivosidenib significantly affect glutathione metabolism in IDH1-mutant cells.**

MetaboAnalyst 6.0 (KEGG) metabolite pathway analysis comparing **A.** untreated and IVO-treated IDH1-mutant cells, or **B.** CMD-treated and CMD-IVO-treated IDH1-mutant cells, or **C.** IVO-treated and CMD-IVO-treated IDH1-mutant cells (20µM IVO for 48 hours, CMD for 17 hours). Top significantly affected pathways are shown with highlighted glutathione, cysteine and methionine metabolism-related pathways.

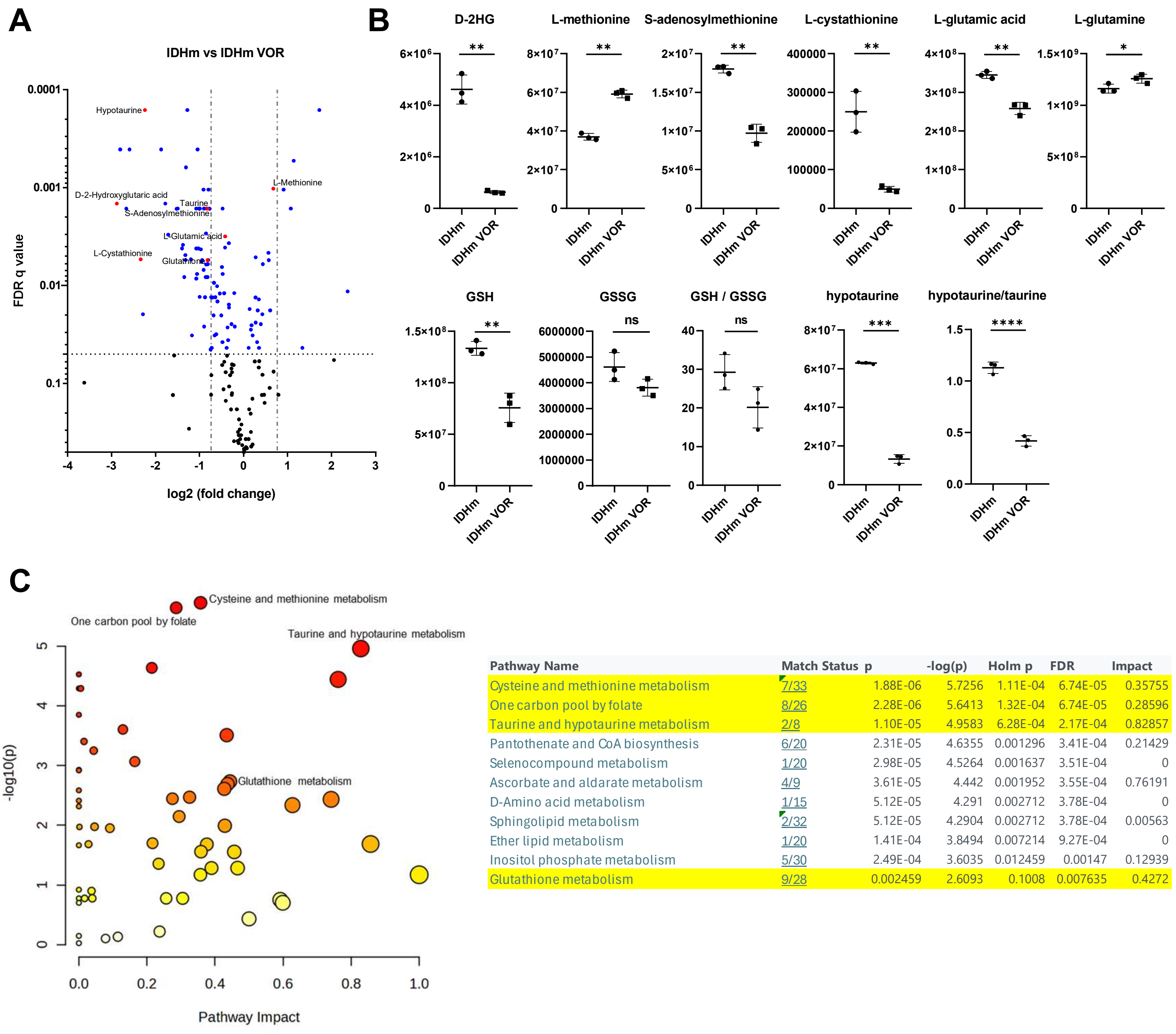


| Pathway Name | Match Status | p | -log(p) | Holm p | FDR | Impact |
|---|---|---|---|---|---|---|
| Cysteine and methionine metabolism | 7/33 | 1.88E-06 | 5.7256 | 1.11E-04 | 6.74E-05 | 0.35755 |
| One carbon pool by folate | 8/26 | 2.28E-06 | 5.6413 | 1.32E-04 | 6.74E-05 | 0.28596 |
| Taurine and hypotaurine metabolism | 2/8 | 1.10E-05 | 4.9583 | 6.28E-04 | 2.17E-04 | 0.82857 |
| Pantothenate and CoA biosynthesis | 6/20 | 2.31E-05 | 4.6355 | 0.001296 | 3.41E-04 | 0.21429 |
| Selenocompound metabolism | 1/20 | 2.98E-05 | 4.5264 | 0.001637 | 3.51E-04 | 0 |
| Ascorbate and aldarate metabolism | 4/9 | 3.61E-05 | 4.442 | 0.001952 | 3.55E-04 | 0.76191 |
| D-Amino acid metabolism | 1/15 | 5.12E-05 | 4.291 | 0.002712 | 3.78E-04 | 0 |
| Sphingolipid metabolism | 2/32 | 5.12E-05 | 4.2904 | 0.002712 | 3.78E-04 | 0.00563 |
| Ether lipid metabolism | 1/20 | 1.41E-04 | 3.8494 | 0.007214 | 9.27E-04 | 0 |
| Inositol phosphate metabolism | 5/30 | 2.49E-04 | 3.6035 | 0.012459 | 0.00147 | 0.12939 |
| Glutathione metabolism | 9/28 | 0.002459 | 2.6093 | 0.1008 | 0.007635 | 0.4272 |

**Supplementary Figure 9. Vorasidenib significantly affects cysteine and methionine metabolism in IDH1-mutant cells.**

**A.** Volcano plot of metabolomics analysis comparing IDH1-mutant (IDHm) untreated cells to IDHm cells treated with 1μM VOR for 48 hours. Blue and red: FDRq<0.05. **B.** Metabolite ratios and metabolites of the methionine cycle, transsulfuration pathway and glutathione metabolism significantly altered by VOR treatment from experiment presented in A. Statistics assessed using t-test. Significance denoted by: *FDRq < 0.05, **FDRq < 0.01, ***FDRq < 0.001, ****FDRq<0.0001, ns: not significant (GSH: Glutathione, GSSG Oxidized glutathione). **C.** MetaboAnalyst 6.0 (KEGG) metabolite pathway analysis comparing IDHm to IDHm VOR-treated cells. Top significantly affected pathways are shown with highlighted glutathione, cysteine and methionine metabolism-related pathways.

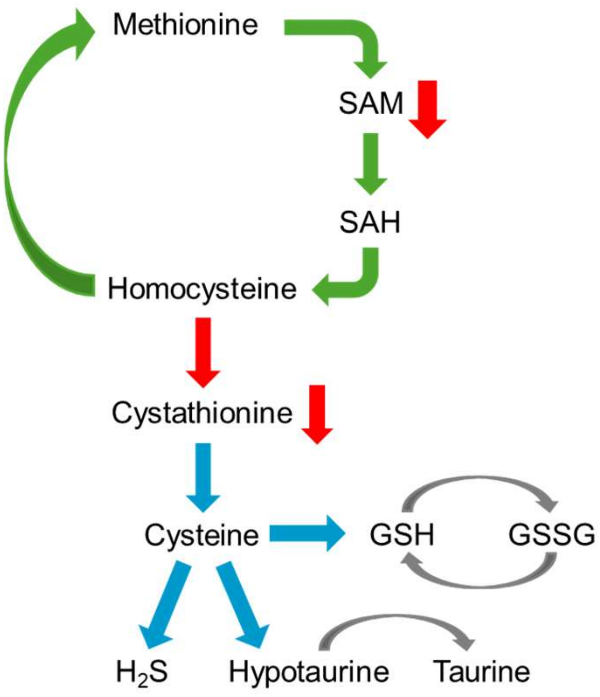


**Supplementary Figure 10. Methionine cycle, transsulfuration and cysteine metabolism.** Diagram of the most significant metabolites of the methionine cycle, transsulfuration and cysteine metabolism. Low S-adenosylmethionine (SAM) levels promote remethylation of homocysteine to methionine and prevent transsulfuration, causing low cystathionine levels. (SAH: S-adenosylhomocysteine, GSH: glutathione, GSSG: oxidized gluathione).

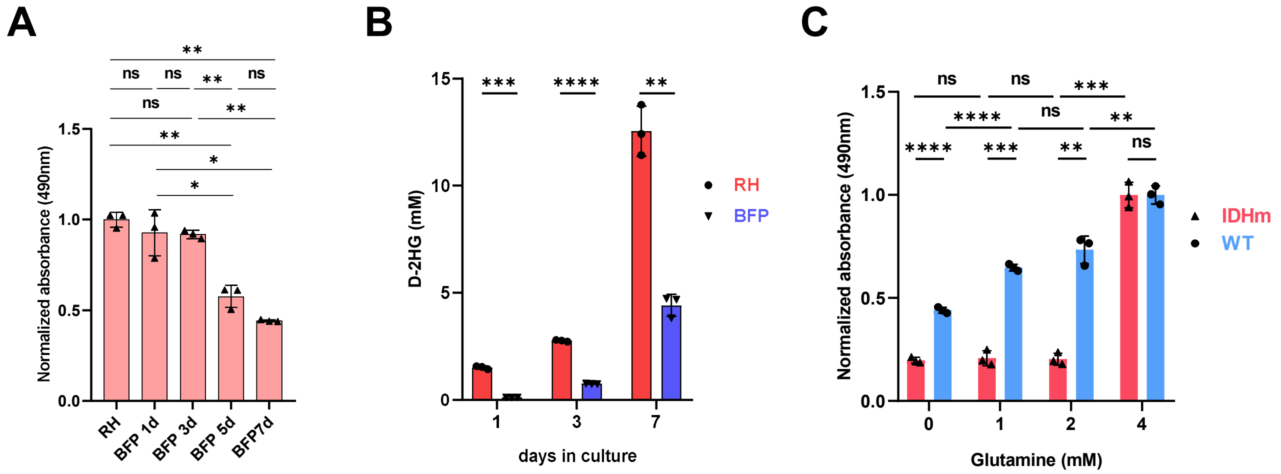


**Supplementary Figure 11. IDH1-mutant cells grown in different conditions express 2-HG *in vitro.***
**A.** MTS viability assay of IDH1-mutant cells grown for 1-7 days in basal media (BFP 1d, 3d, 5d, 7d) compared to IDH1-mutant cells grown in enriched media for 3 days (RH). **B.** 2-HG measured in IDH1-mutant cells grown in parallel in either basal (BFP) or enriched (RH) media. **C.** MTS viability assay of IDH1-mutant and IDH1-wildtype cells grown for 2 days in basal media containing 25mM glucose and different concentrations of glutamine (0, 1, 2, 3, 4 mM). Data is presented as mean ± SD. Statistics assessed using t-test. Significance denoted by: *$p < 0.05$, **$p < 0.01$, ***$p < 0.001$, ****$p<0.0001$, ns: not significant.

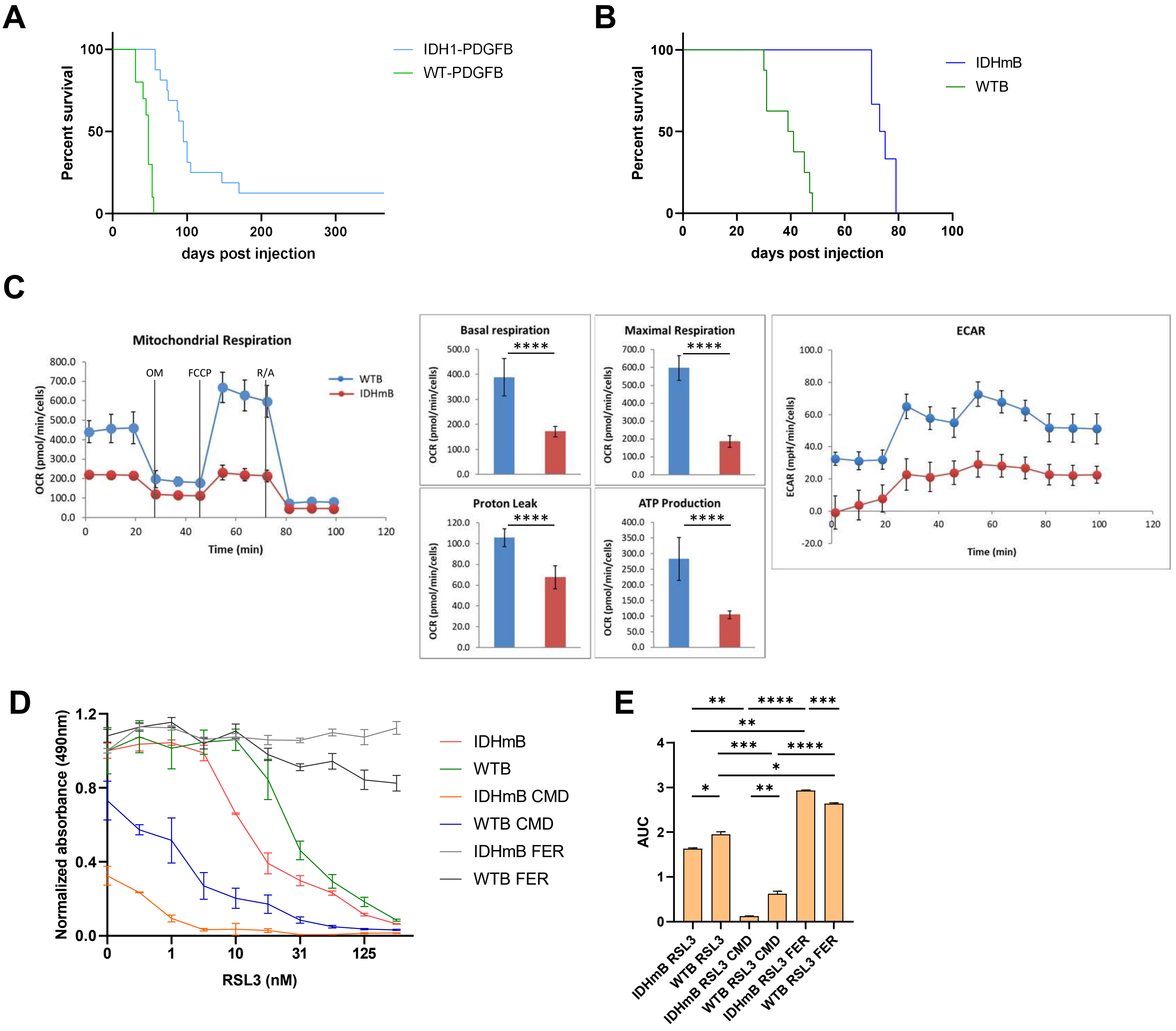


**Supplementary Figure 12. IDH1-mutant, PDGFB-expressing glioma cells induce tumorigenesis, are sensitive to oxidative stress and ferroptosis.**

**A.** Survival curves after stereotactic intracranial retrovirus injections in neonatal (p4) mice. Idh1(R132H)$^{KI/+}$; p53$^{fl/fl}$; RiboTag and +/+; p53$^{fl/fl}$; RiboTag mice injected with PDGFB-IRES-CRE retrovirus (IDH1-PDGFB and WT-PDGFB). Median survival: IDH1-PDGFB 95dpi, WT-PDGFB 48dpi. Log-Rank (Mantel-Cox) test: $p<0.0001$****. **B.** Survival curves after orthotopic injections of IDH1-mutant PDGFB-expressing cells (IDHmB) and IDH1-wildtype PDGFB-expressing (WTB) cells into adult B6 mice. Median survival: IDHmB 74dpi, WTB 40dpi. Log-Rank (Mantel-Cox) test: $p=0.0003$***. **C.** Seahorse Mitochondrial stress test of IDH1-wildtype (WTB: blue) and IDH1-mutant cells (IDHmB: red) in BFP media (n = 6 per group). OM: oligomycin, FCCP: Carbonyl cyanide-4 (trifluoromethoxy) phenylhydrazone, R/A: rotenone and antimycin A. Basal respiration, maximal respiration, ATP production and proton leak values were calculated and normalized. Extracellular acidification rate (ECAR) is also presented. **D.** Dose response curves and **E.** AUC after treatment with RSL3 (0, 0.5, 1, 5, 10, 15, 31, 62, 125, 250 nM) in control or CMD conditions, with or without 2μM Ferrostatin-1 (IC50 WTB 36.7, IDHmB 10.1, WTB-CMD 9.601, IDHmB-CMD 0.4555). Data presented as normalized mean ± SD. Statistics assessed using t-test. Significance denoted by: *$p < 0.05$, **$p < 0.01$, ***$p < 0.001$, ****$p<0.0001$.
